# Analysing diffusion-limited processes in a cylinder using pair-correlation functions

**DOI:** 10.1101/2025.02.19.639196

**Authors:** Benjamin James Binder

## Abstract

Diffusion-limited processes (DLP) are found in various physical, biological, and engineering systems, yet their quantification within complex spatial domains remains a challenge. In this study, we develop novel one-dimensional non-periodic and periodic pair-correlation functions (PCF) to assess the spatial patterns of DLP within a cylindrical domain. By refining previous PCF formulations, we introduce an efficient binning-based approach that significantly reduces computational costs, making the method feasible for large-scale simulations. Our analysis provides a comprehensive examination of PCF variability, distinguishing between global deviations from complete spatial randomness state and sampling-induced variation. An off-lattice agent-based model is implemented, successfully reproducing self-organized patterns reminiscent of classical DLP studies and aligning with fractal-like aggregation behaviours. We demonstrate the utility of periodic PCFs in capturing key spatial correlations in DLP, particularly in azimuthal and Cartesian projections, while highlighting the conditions under which non-periodic PCFs remain preferable. Our findings underscore the potential of PCFs as robust summary statistics for complex spatial models, with applications ranging from microbial colony formation and blood clotting dynamics to image analysis and classification algorithms.

## I. INTRODUCTION

Diffusion-limited processes (DLP) refer to phenomena where the transport of particles (e.g. molecules, atoms, or cells) via diffusion is the primary determinant of the dynamics, efficiency, or rate of an interaction or transformation [1, 2]. These processes occur in physical systems when diffusion is significantly slower than other mechanisms, such as reaction kinetics, material deposition, or biological growth, thereby becoming the rate-limiting step in the system’s overall behaviour. DLP are widely observed across various physical systems and examples include crystal and dendrite growth [3], electrodeposition [4], microbial colony and biofilm formation [5], the Saffman–Taylor problem in fluid mechanics [6], and the development of lightning strikes [7]. Additional examples of DLP within a cylindrical domain are illustrated in Fig. 1.

**FIG. 1.**
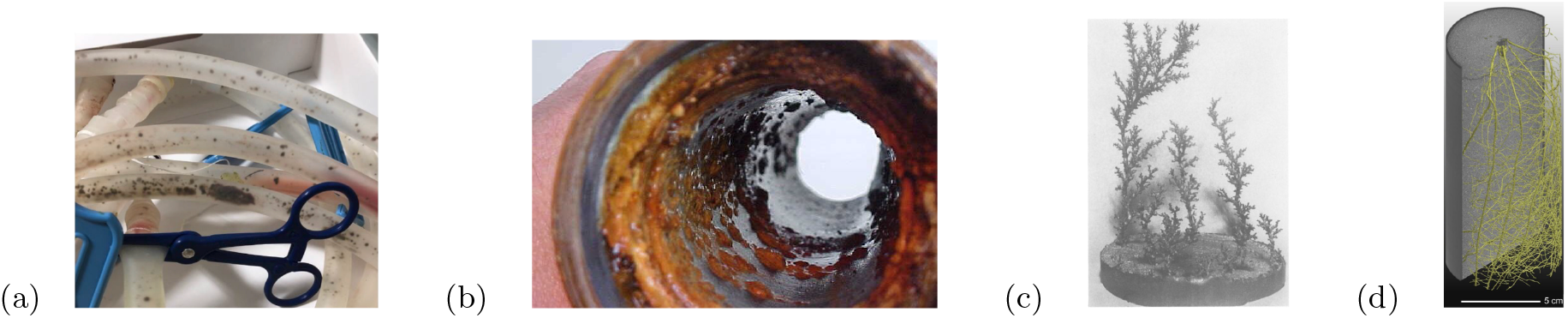
Physical examples of diffusion-limited processes within a cylindrical domain. (a) Microbial colonies of black mold in bilateral chest tubing [12]. (b) Macro-biofilm growth on waste water pipe [13]. (c) Dissolution of plaster of Paris [14]. (d) Rendered 3D image of maize root growth [15].

Quantifying spatial patterns of DLP is essential for understanding how systems evolve under constraints imposed by diffusion. For example, diffusion-limited growth (DLG) occurs in microbial colonies when nutrient availability restricts growth, leading to non-uniform morphologies. Bacterial colonies grown in Petri dishes exhibit hallmark DLG features, including branch screening, colony repulsion, and directed growth toward nutrient sources [5]. These characteristics have been successfully reproduced using lattice-based agent models [8]. Similarly, yeast colonies grown under nutrient-limited conditions exhibit highly non-uniform spatial patterning [9]. However, Tronnolone et al. [10, 11] demonstrated that yeast colonies lack the DLG trait of directed growth toward nutrient sources. Instead, their non-uniform morphologies were attributed to cell proliferation mechanisms rather than diffusion-limited effects. This example shows the need for spatial metrics capable of characterising non-uniform patterning to effectively evaluate the role of DLP in physical systems.

Another example of diffusion-limited processes (DLP) can be found in the seminal work of Witten and Sander [16, 17], where a lattice-based agent model was developed to simulate diffusion-limited aggregation (DLA) in the *xy*-plane. To quantify the patterning observed in their simulations, they used the density-density correlation function which measures how the density at one point in space is correlated with the density at another point. An alternative measure of spatial correlation is the Euclidean pair correlation function (PCF), also known as the radial distribution function [18].

The Euclidean PCF describes how objects or points are spatially distributed relative to each other in a system [19, 20]. It measures the relative probability of finding two objects separated by a Euclidean distance *r* compared to a completely random distribution with the same density. The PCF is defined as

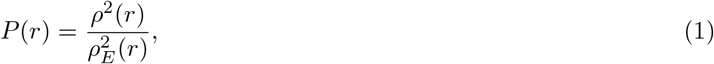

where *ρ*^2^(*r*) is the observed density of pairs at distance *r* in the system and 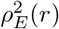 is the expected pair density at distance *r* for a uniform random distribution of objects within the spatial domain. A system with *P* (*r*) = 1 at all distances *r* indicates no clustering or regular patterning, often referred to as a state of complete spatial randomness (CSR) [21–24]. When *P* (*r*) *>* 1 at certain distances, there is an increased probability of finding objects close together, indicating clustering or aggregation. Conversely, when *P* (*r*) *<* 1, there is a reduced likelihood of observing objects, suggesting regular spacing or segregation in the system.

The Euclidean PCF is widely applied to assess patterning in unconstrained spatial domains (e.g. the *xy*-plane). However, it has limitations in systems where Euclidean distance is not the most appropriate measure, such as lattice-based models at relatively short to moderate distances. Binder and Simpson [25] partially addressed this limitation by developing a one-dimensional Cartesian PCF for projected patterns along either the *x* or *y* axis in square-lattice simulations. Subsequently, Gavin et al. [26] introduced the taxicab distance as a metric in their formulation of the Manhattan PCF, extending its application to square lattices in two and three dimensions, as well as to non-square lattices. Binder et al. [9] implemented a one-dimensional azimuthal PCF in cylindrical coordinates to quantify dimorphic transitions in the growth modes of filamentous yeast colonies. Similarly, Dini et al. [27] employed one-dimensional polar, azimuthal, and radial PCFs in spherical coordinates to delineate the necrotic zone boundary in tumuour spheroids. Moreover, PCFs with alternative distance metrics beyond the Euclidean framework have been explored for multi-species spatial patterns, domains with obstacles, and high-dimensional spaces [28–30].

A significant limitation in all the aforementioned studies is the efficient evaluation of the pair correlation function (PCF) when the number of objects (e.g. agents, cells, molecules), *n*, in the domain of interest is large. The PCF calculation requires 𝒪 (*n*^2^) evaluations of the distances between all pairwise combinations of objects in the domain, regardless of the distance metric used. Computationally this becomes prohibitive as *n* increases. One approach to mitigate this issue is to take a random sample of *n*_*s*_ ≪ *n* objects from the domain, calculate the PCF for each sample, and then compute an average PCF to estimate the expected PCF of the process [9, 25, 31, 32]. However, this method has its own shortcomings, particularly when considering multiple realisations or experiments of the same process. In such cases, an average of the average PCFs from the samples is often used to estimate the PCF of the underlying process. This approach makes it challenging to analyse the variability of the underlying process using straightforward and interpretable statistical techniques, as it complicates the application of standard methods for assessing uncertainty or variability in the results.

The aim of this work is to develop new one-dimensional PCFs for projected patterns in the azimuthal, Cartesian, and radial directions to efficiently characterise DLP spatial patterns with large numbers of objects constrained within a cylindrical domain. This is achieved by binning the projected data into *N≪ n* bins and evaluating the pairwise combinations of the counts in these *N* bins instead of the pairwise combinations of the *n* objects in the domain, significantly reducing the computation time required to estimate the PCFs. An obvious advantage of this approach is that the number of pairwise combination evaluations does not increase for a fixed number of bins, even as the number of objects in the domain grows. This allows direct sampling of DLP processes without the need to sample individual realisations or experiments, thereby reducing the layers of variation to analyse when estimating the expected PCF for a given process.

In our reformulation of the estimate of the one-dimensional PCF for projected patterns, we derive expressions (or normalisation terms) for the expected number of counts for each pairwise bin combination in a cylindrical domain populated uniformly at random (i.e. terms analogous to 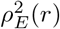 Sin Eq. (1)). Furthermore, we develop a weighted mean value of the PCF that always sums to unity across all distances, enabling straightforward analysis of variability for a given DLP process. Importantly, we demonstrate that the total variability,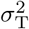, can be expressed as the sum of two components: the variability within the sample of the process, *s*^2^, and the deviation of the process from the CSR state, 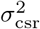. This allows our PCFs to identify departures from the CSR state at specific distances (when the PCF is not equal to unity) and to quantify the overall deviation from the CSR state. Additionally, we show that the discrete cosine transform [32, 33] of the PCF can be used to express 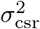 in terms of the transform coefficients, and examining the power spectrum enables the identification of characteristic length scales in DLP processes.

In the next section, we define an off-lattice model for DLP within a cylindrical domain and evaluate the effectiveness of our PCFs in quantifying the spatial patterns generated by the model. Our results demonstrate that computationally efficient PCFs can accurately characterise a wide range of spatial patterns within the cylindrical domain framework.

## II. FORMULATION

In this section, we define an off-lattice model for DLP and derive the one-dimensional PCFs for projected patterns in the azimuthal, Cartesian, and radial directions.

### A. Agent-based model

We consider a discrete off-lattice model to simulate DLP within a cylindrical domain, (*r, θ, z*). Within the domain, there are a total of *J* species, and at time *t*, the total number of agents in the *j*th species is denoted by *n*^*j*^(*t*). In the Cartesian framework, the location of the *i*th agent of the *j*th species is described by

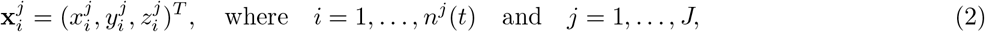

with corresponding position vectors in the cylindrical coordinate system given by

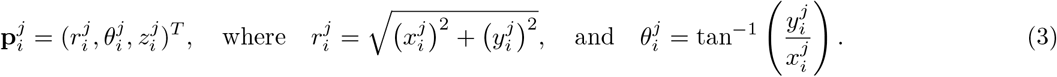

Note that this location notation does not apply in the degenerate case(s) where there are no agents in the *j*th species at time *t*, i.e. when *i* = *n*^*j*^(*t*) = 0.

In the simulations, we use a finite-size domain with 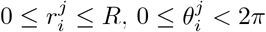 and 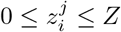.

#### 1 Motility and transport events

The movement of agents within the cylindrical domain can be broken down into two components. The first component is a random walk, where each agent is given the opportunity to move during a time step, *δt*. The second component is a variable drift in the *z* direction. During a time step, the position of each agent attempts to move to

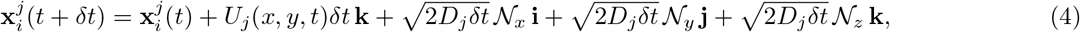

where *D*_*j*_ is the diffusion coefficient and *U*_*j*_ is the drift for the *j*th species, [34]. The terms 𝒩_*x*_, 𝒩_*y*_, and 𝒩_*z*_ are independent Gaussian random variables, each with zero mean and unit variance. The movement event is aborted if the agent’s target location lies outside the cylindrical domain, i.e. if 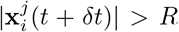. Collision events between individual agents during a time step are not considered in the model due to the prohibitive computational cost of simulating agent interactions within the domain.

#### 2 Diffusion-limited processes

During each time step of the algorithm, diffusion-limited processes allow for a decrease in the number of agents in one species accompanied by an increase in the number of agents in another species. For example, one species might represent nutrients, which can be consumed to promote growth in a second species representing biological cells. From a modelling perspective, this is equivalent to an aggregation process, where the first species represents a quantity of interest suspended in a fluid or solute, and the second species represents the precipitate of the same material forming a solid aggregate.

The removal of an agent in the first species and the associated addition of a new agent in the second species can be implemented as follows. For simplicity, we describe a biological growth process with two species, where one nutrient agent is sufficient to produce one biological cell. Let the nutrient and cell agents be denoted by 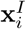 and 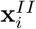 respectively. We consider the birth of a new daughter cell adjacent to an existing mother cell located at 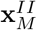. The mother cell searches for nutrient agents within its local environment, defined by a search radius *δr*. This search identifies the set of nutrient agent locations as

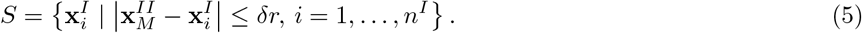

A member of the set *S* is selected at random and is chosen as the location of the new daughter cell, with the nutrient agent at this location being removed from the domain.

The basic rule can be readily modified to accommodate additional constraints and processes. For example:

- A rate or probability, 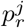, can be introduced to model the likelihood that biological cells absorb or consume nutrient agents (*j* = *I*), adding stochasticity to the absorption process.
- If more than one nutrient agent is required to produce a biological cell, the process can be adapted to remove nutrient agents within the mother cell’s search radius until the required number is reached. The location of the last nutrient agent removed can then be chosen as the location of the new daughter cell.

A simulation of the model with two species is shown in Fig. 2. The first species, *I*, is motile and undergoes DLP, transforming its agents into species *II*. The second species, *II*, is immotile and does not degrade, decrease in number, or undergo any further DLP into a third species. Initially, *n*^*I*^ (0) = 6000 agents of species *I* are distributed uniformly at random throughout the cylindrical domain, with one agent of species *II* (*n*^*II*^ (0) = 1) placed at (0, 0, *Z*) as a seed to enable DLP. The simulation is terminated when *n*^*I*^ = 0 and *n*^*II*^ = 6001.

**FIG. 2.**
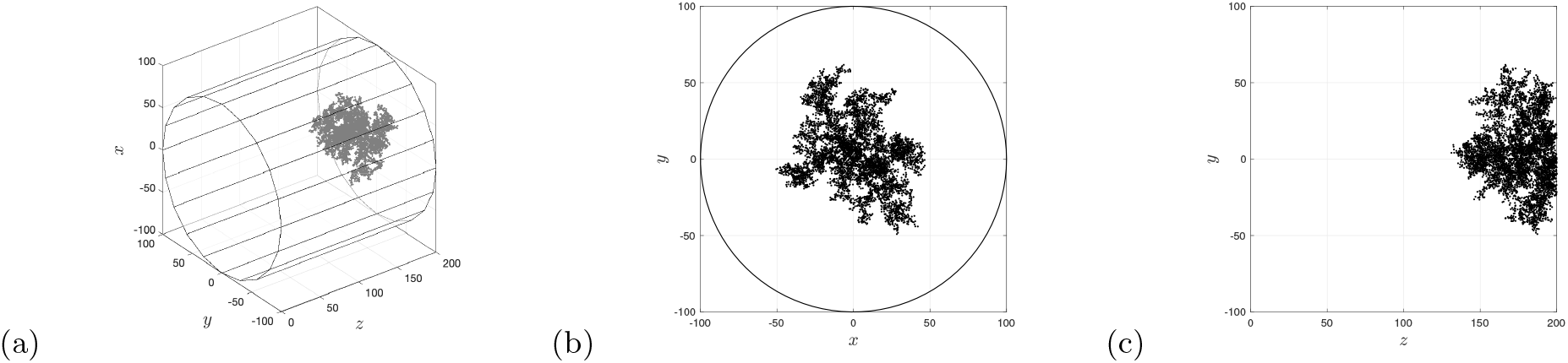
Diffusion-limited processes within a close-ended cylinder, *R* = 100, *Z* = 200, *D*_*I*_ = 1, *D*_*II*_ = *U*_*I*_ = *U*_*II*_ = 0, *δt* = 1, *δr* = 2. Initially *n*^*I*^ (0) = 6000 agents of species *I* were distributed uniformly at random throughout the cylinder with a single agent from species *II* located at (0, 0, *Z*) with *n*^*II*^ (0) = 1. The simulations shown in (a)–(c) were terminated when all the agents in species *I* had transformed into species *II*. (a) Simulation in cylindrical domain, *t* = 133, 160. (b) Projection in *rθ*-plane. (c) Projection in *zy*-plane (b).

Next, we derive one-dimensional PCFs to quantify the spatial patterning in cylindrical domains.

### B. Pair-correlation functions

We develop both a non-periodic and periodic [25, 35] one-dimensional pair-correlation function (PCF) to assess the spatial patterning of *n*^*j*^ agents with position vectors 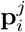 in a cylindrical domain (*r, θ, z*). The first step in the derivation is to define equal-size volumes of three-dimensional space and their corresponding projections onto the one-dimensional interval [0, *L*] for each of the three orthogonal directions, *r, θ*, and *z*. The projected positions of the agents will then reside on the intervals [0, *L*], with *L* = *R*, 2*π, Z*, for agents located within a cylinder of radius *R* and length *Z*. Equal-size bin widths on the intervals [0, *L*] in the *θ* and *z* directions, with *L* = 2*π* and *L* = *Z*, correspond to equal-size volumes in three-dimensional space (i.e. equal-volume segments and equal-volume smaller-length cylinders). However, special care is required in the radial direction, as equal-size bin widths on [0, *R*] do not correspond to equal volumes in three-dimensional space. To address this, we define discrete distances or *N* equal-size bins for the *θ* and *z* projections on [0, *L* = 2*π, Z*] as

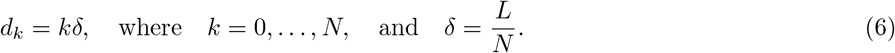

For the radial case, we replace (6) with *N* variable bin widths on the interval [0, *L* = *R*]

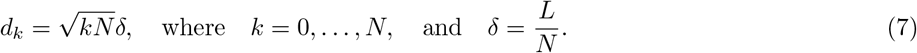

The variable bin widths in (7) correspond to a small central cylinder of radius *δ* and *N ™* 1 concentric annular cylinders, all with the same equal-size volume *v* = *πδ*^2^*Z* in three-dimensional space. (In the Appendix, we provide the variable bin widths for the spherical coordinate system, demonstrating that the PCFs developed in this work can be applied to other curvilinear coordinate systems.)

The corresponding sets of the projected agent positions 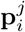 onto each of the orthogonal unit vectors **ê**_**r**_, **ê**_*θ*_, and **ê**_**z**_ of the cylindrical coordinate system are given by

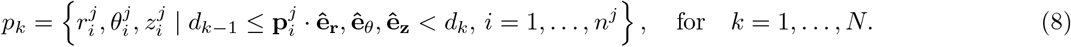

The number of elements in each set, or the bin-count at each dimensionless non-zero discrete distance, *k*, can then be defined as

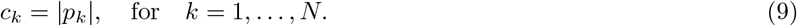

As previously mentioned in the Introduction, an important difference in the derivation presented here is the consideration of pair-distance bin-counts instead of pair-distance of individuals (e.g. agents) as found in the authors previous work. This significantly reduces computation of the PCF because the number of paired-bin combinations remains constant for a fixed number of bins, even when the number of agents increases in the calculation of the PCF (see Appendix). Hence, for dimensionless non-zero pair-distances, we define two sets of non-periodic and periodic pair-distance bin-counts as

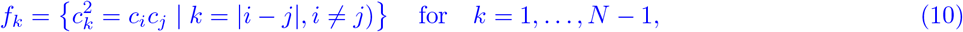

and

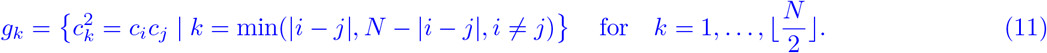

respectively. The zero pair-distance counts are simply given by the total number of pairs of agents in each of the *N* bins,

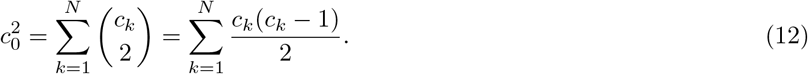

The PCFs are defined by scaling the sum of the elements in the families of paired-bin counts (10)–(12) such that a domain populated uniformly at random is expected to return a unit-valued PCF at all pair distances. The non-periodic and periodic PCFs are given by

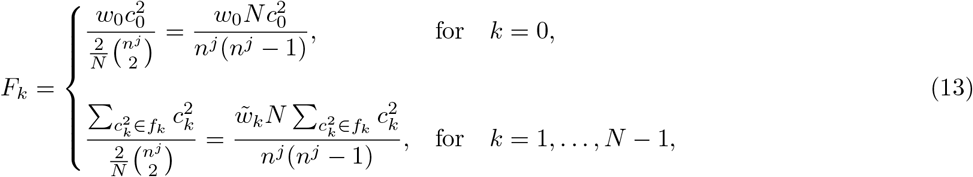

and

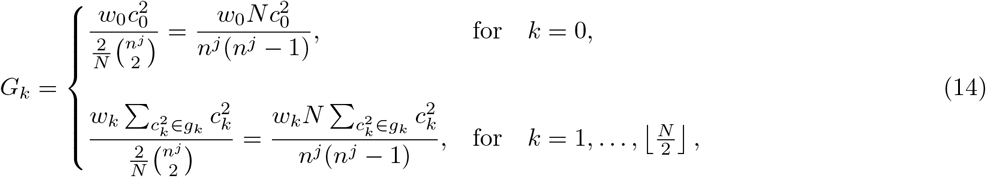

where 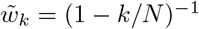, *w*_0_ = 2 and *w*_*N/*2_ = 2 for an even number of bins; otherwise, *w*_*k*_ = 1. We remark that the term 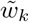 in the non-periodic formulation is similar to that found in Binder and Simpson’s original work. The pairwise bin approach here shares some commonality with their formulation for pairwise individuals on a square lattice [25].The combinatorial term in both PCFs essentially represents the probability of selecting two agents (or bins when scaled by *N*) in the spatial domain.

The PCFs defined in (13)–(14) can be used to assess the spatial patterning in the Cartesian, azimuthal, and radial directions. Evaluation of the PCFs for a cylindrical domain that has been populated uniformly at random with 2000 agents is shown in Fig. 3. At short to moderate distances, both the periodic (red) and non-periodic (blue) PCFs fluctuate closely around unity, as expected. The larger fluctuations in the non-periodic PCF at longer distances are due to the division by small numbers in the metric.

**FIG. 3.**
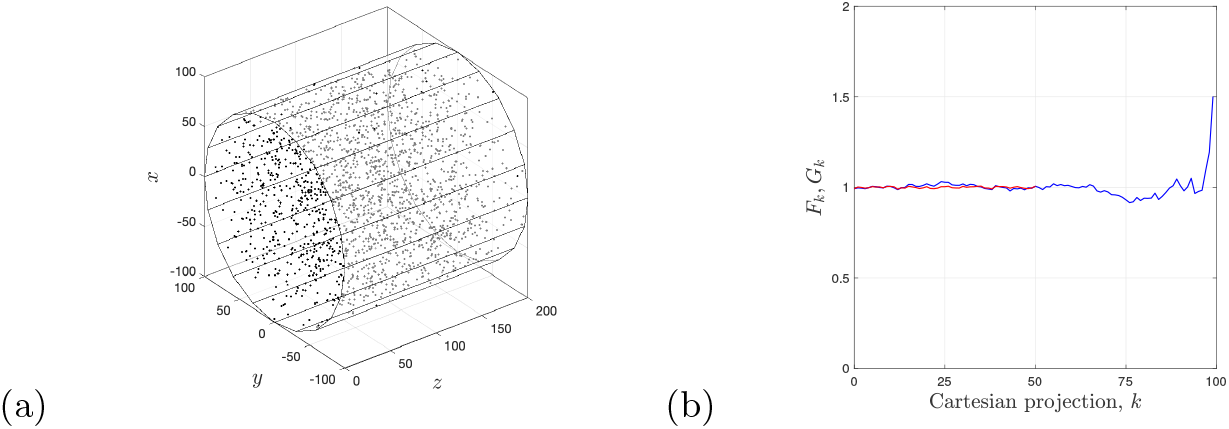
Cylinder populated uniformly at random with *n*^*I*^ = 2000 agents, *R* = 100, *Z* = 200. (a) Populated domain. (b) Non-periodic (blue curves) and periodic (red curves) PCFs, *F*_*k*_ and *G*_*k*_ for the projection in *z*-direction. Similar results are found for the *θ, r*-directions (not shown).

#### 1. Cartesian, azimuthal and radial metric

We modify the metrics of Binder et al. [9] and scale histograms of the projected data for comparison with the PCFs. For each of the three projections, [0, *L* = *Z*, 2*π, R*], we define *N* bins with equal-size bin widths, *δ* = *L/N*, and corresponding counts, *C*_*k*_. A scaled histogram of the counts is then given by

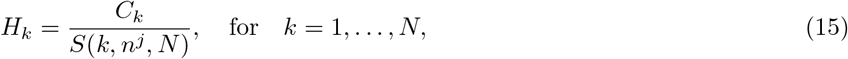

where *S*(*k, n*^*j*^, *N*) is chosen to return an expected unit-valued metric at all distances for a domain populated uniformly at random. For the Cartesian and angular projections, the bins are equal in size, and the scaling is simply *S* = *n*^*j*^*/N* (with *C*_*k*_ = *c*_*k*_). In the radial case, however, the bins are not equal in size, and the scaling is

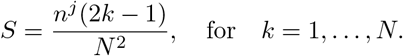

#### 2. Average metrics

We also calculate the average of the metrics 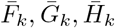 from *M* realizations. For example, the average periodic pair-correlation function is defined as

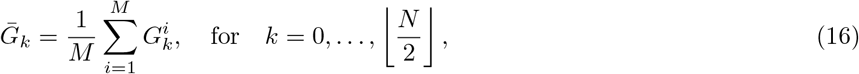

with variance

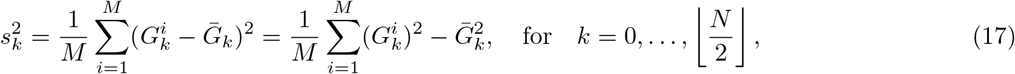

where 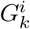 is the periodic PCF for the *i*th realization. Similar expressions hold for the non-periodic PCF and scaled histogram metrics.

To examine the accuracy of our derivations, we compute the average PCFs from *M* = 1000 realizations of a cylinder populated uniformly at random with *n*^*I*^ = 2000 agents (see Fig. 4). The results illustrate that the average of both the non-periodic and periodic PCFs approach unity at all valid distances, as expected. In addition, we plot two standard deviations (dotted curves) of the average PCF. Consequently, we expect the PCF from a single realization to lie between the two dotted curves with 95% confidence. Consistent with the authors’ previous work, we find that the non-periodic PCF exhibits greater uncertainty or variability than the periodic PCF, particularly at large distances. However, both PCFs demonstrate a much higher degree of accuracy compared to the average scaled histogram metric, again as expected.

**FIG. 4.**
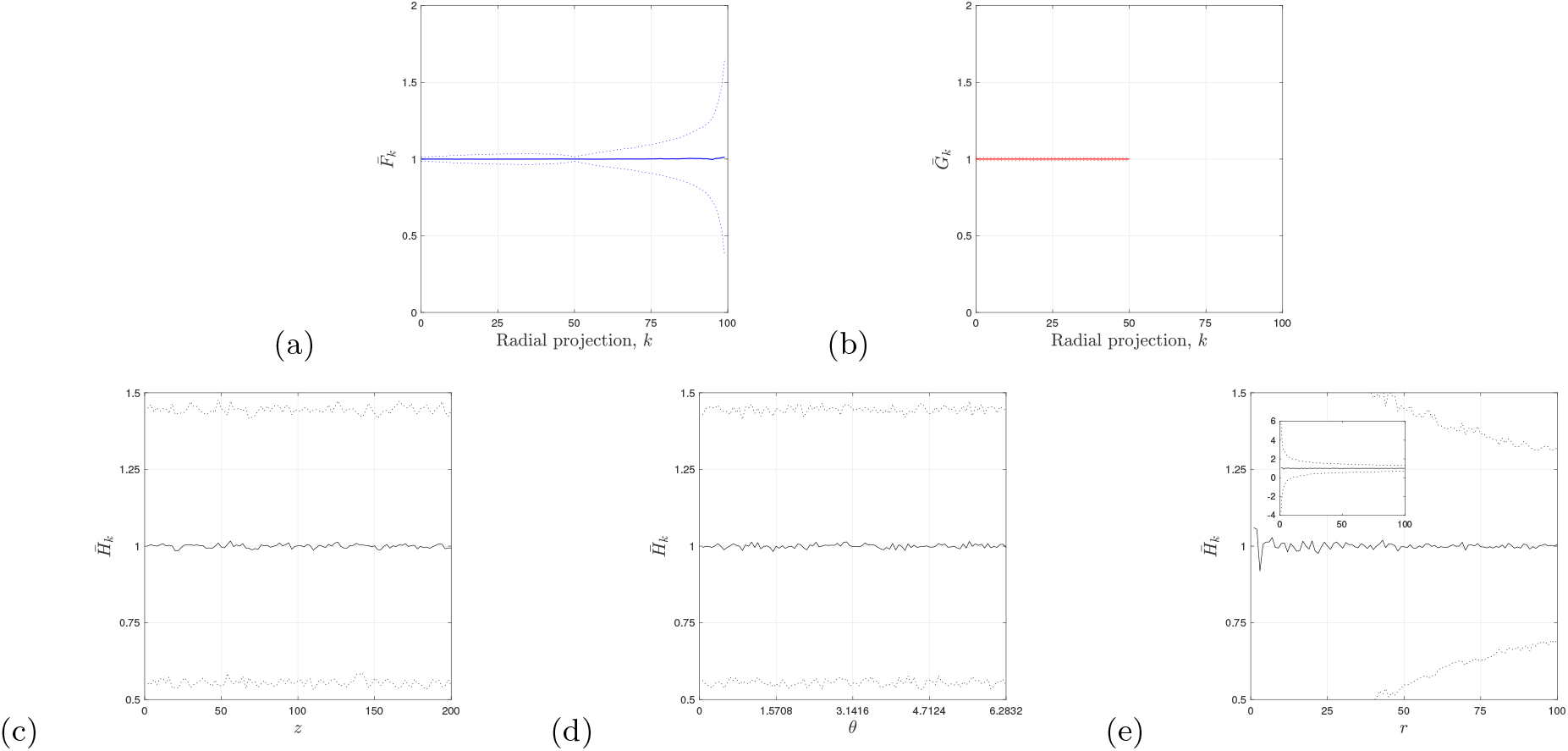
Cylinder populated uniformly at random with *n*^*I*^ = 2000 agents, *R* = 100, *Z* = 200, *N* = 100. Average of spatial metrics calculated from *M* = 1000 realisations, solid curves. The dotted curves are for 2 standard deviations from the mean. and (b) Average non-periodic (blue curves) and periodic (red curves) PCFs, 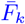 and 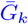, for the radial projection. Similar results are found for the *θ, r* projections. (c)–(e) Scaled histogram metric (black curves), 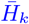. (c) Projection in *z*-direction. (d) Projection in *θ*-direction. (e) Projection in *r*-direction.

Having demonstrated that the PCFs can accurately predict when patterns are close to the CSR state in a cylindrical domain, we now investigate non-trivial patterns generated by the DLP model. In the following analysis, we describe the projected patterns at *distances* along the direction of projection, where PCF values greater than one indicate aggregation and values less than one indicate segregation. Patterns are considered close to the CSR state when the PCF values are near unity, and note that this terminology refers solely to the projection being analysed and does not imply that the overall pattern is at the CSR state. Rather, the CSR state is used as a reference point to compare our results.

It is important to discuss the interpretation of *distance* for each of the projections. Recall the dimensionless nonzero distances *k* = 1, …, *N*. These dimensionless distances correspond to an ordered list of equal volumes in 3D space, regardless of the coordinate system or projection. For example, a dimensionless distance of *k* = 2 refers to equal volumes separated by one equal volume in 3D space. When the projection has equal or constant bin widths, as in the Cartesian and azimuthal projections, the dimensionless distances correspond directly to the physical distances *d*_*k*_ in (6). However, this is not the case when variable bin widths are present, as in the radial projection (7). Therefore, PCFs in the Cartesian and azimuthal projections are plotted as functions of the physical distances, *d*_*k*_, whereas the radial PCFs are plotted as functions of the dimensionless distance, *k*.

## III. RESULTS

We begin the discussion by examining the spatial patterning of DLP in the two-dimensional (*r, θ*)-plane using the PCFs. Initially, *n*^*I*^ (0) = 2000, 4000, 6000 agents of species *I* are distributed uniformly at random throughout a circle, with a single agent from species *II* located at the origin (*n*^*II*^ (0) = 1). The simulations are terminated when all the agents in species *I* have transformed into species *II*, as shown in the first column of Fig. 5.

**FIG. 5.**
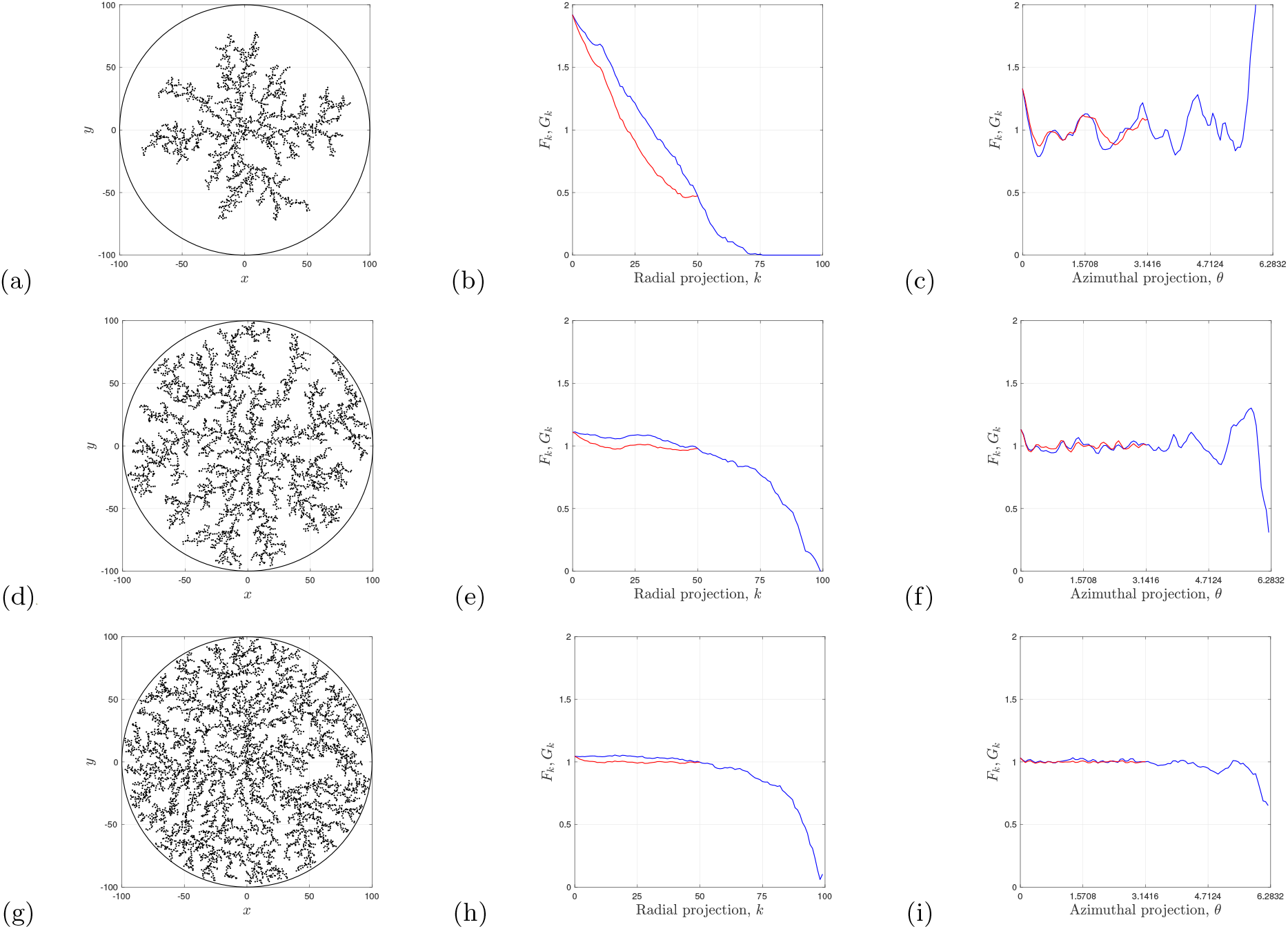
Diffusion-limited processes within a circular domain, *R* = 100, *D*_*I*_ = 1, *D*_*II*_ = *U*_*I*_ = *U*_*II*_ = 0, *δt* = 1, *δr* = 2. First column. Initially, *n*^*I*^ (0) = 2000, 4000, 6000 agents of species *I* were distributed uniformly at random throughout the circle with a single agent from species *II* located at the origin with *n*^*II*^ (0) = 1. The simulations were terminated when all the agents in species *I* had transformed into species *II*. Non-periodic (blue) and periodic (red) PCFs, and scaled histogram metrics (grey), *N* = 100. Second column. Projection in *r*-direction. Third column. Projection in *θ*-direction. (a)–(c) *t* = 4, 381. (d)–(f) *t* = 1, 372. (g)-(i) *t* = 478.

The 2D results from our off-lattice DLP model closely resemble the (square) on-lattice model of Witten and Sander [16, 17], exhibiting the characteristic branch screening patterns that prevent the central region of the domain from becoming fully occupied with agents (see first column, Fig. 5). The patterning in the radial direction is quantified with the PCFs, as shown in the second column of Fig. 5. In panel (b), both the non-periodic and periodic PCFs decrease as the distance increases, which is expected because the density decreases with increasing radial distance from the origin. However, as the number of agents increases in the simulations (e, h), the decrease in the radial PCFs with distance is much less pronounced. This behaviour arises because the density is nearly constant at short to moderate distances in the radial direction. In contrast, the PCFs for the *θ*-direction (third column, Fig. 5) fluctuate around unity at most distances, making it more challenging to assess the pattern without examining the average PCFs for the process.

The average behaviour of the results under the same conditions as those presented in the first row of Fig. 5 is considered next. In the radial direction, the PCFs from a single realisation (dotted curves) closely follow the average PCFs from *M* = 1000 realisations (solid curves), as shown in Fig. 6(a, b). The description of the average behaviour in the radial direction is, therefore, more or less the same as previously observed for a single realisation of the model.

When examining results in the *θ*-direction, we find that the fluctuations observed in a single PCF signal are smoothed out in the average PCF signal [Fig. 6(d, e)]. The average PCF signal is at unity for moderate distances, indicating that the patterning is close to the CSR state at those distances in the *θ*-direction. On closer examination of the average non-periodic PCF signal, we observe near symmetry about *θ* = *π*, with aggregation at very short and very large distances [Fig. 6(d)]. This behaviour can be explained as follows. At very short distances, aggregation occurs because agents of species *I* transform into agents of species *II* very close to an existing agent of species *II*. Aggregation at very large distances is due to the 2*π* periodicity in the azimuthal direction. Agents that are close together can have angles close to either zero or 2*π*, giving the somewhat misleading signal of aggregation at very large distances. This counter intuitive result of the non-periodic PCF is not an issue in periodic PCF signal where we only observe aggregation at very short distances [Fig. 6(e)].

**FIG. 6.**
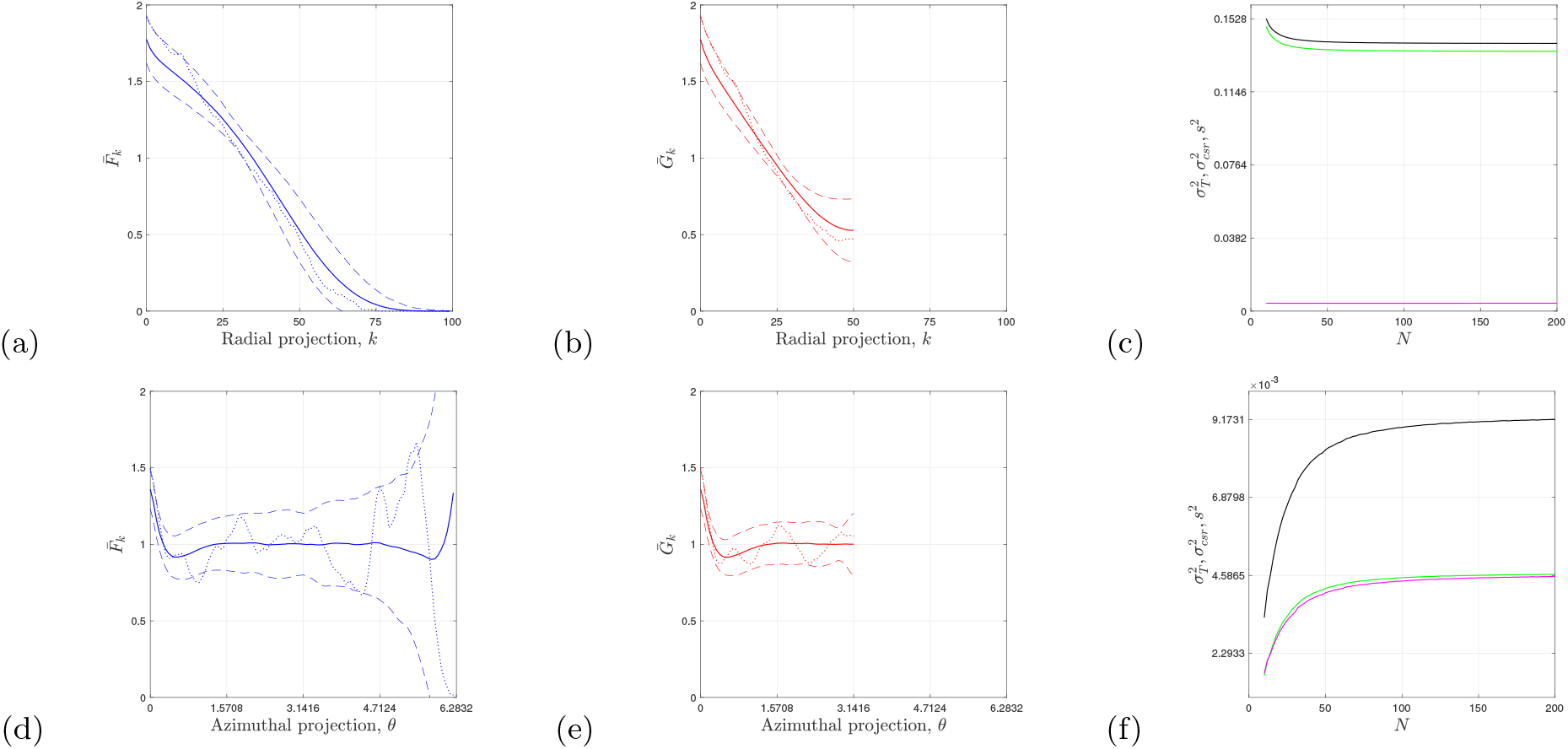
Diffusion-limited processes within a circular domain, *R* = 100, *D*_*I*_ = 1, *D*_*II*_ = *U*_*I*_ = *U*_*II*_ = 0, *δt* = 1, *δr* = 2. Initially, *n*^*I*^ (0) = 2000 agents of species *I* were distributed uniformly at random throughout the circle with a single agent from species *II* located at the origin with *n*^*II*^ (0) = 1. The simulations were terminated when all the agents in species *I* had transformed into species *II*. (a), (b), (d), (e) Average of PCFs calculated from *M* = 1000 realisations, solid curves. The dashed curves are for 2 standard deviations from the mean. The dotted curves are for a single realisation. Non-periodic (blue) and periodic (red) PCFs, *N* = 100. (c) and (f) Analysis of variance from *M* = 1000 realisations, 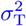 (black), 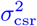 (green), *s*^2^ (magenta). (a)-(c) Projection in *r*-direction. (d)-(f) Projection in *θ*-direction.

The results for the average PCFs under the remaining two sets of conditions, corresponding to the second and third rows of Fig. 5, are presented in the first two columns of Figs. 7 and 8. The qualitative behaviour remains consistent with that observed in Fig. 6. However, there are notable quantitative differences when comparing the three sets of average results for *n*^*I*^ (0) = 2000, 4000, 6000. These differences highlight the ability of the PCFs to distinguish between the three distinct sets of conditions.

**FIG. 7.**
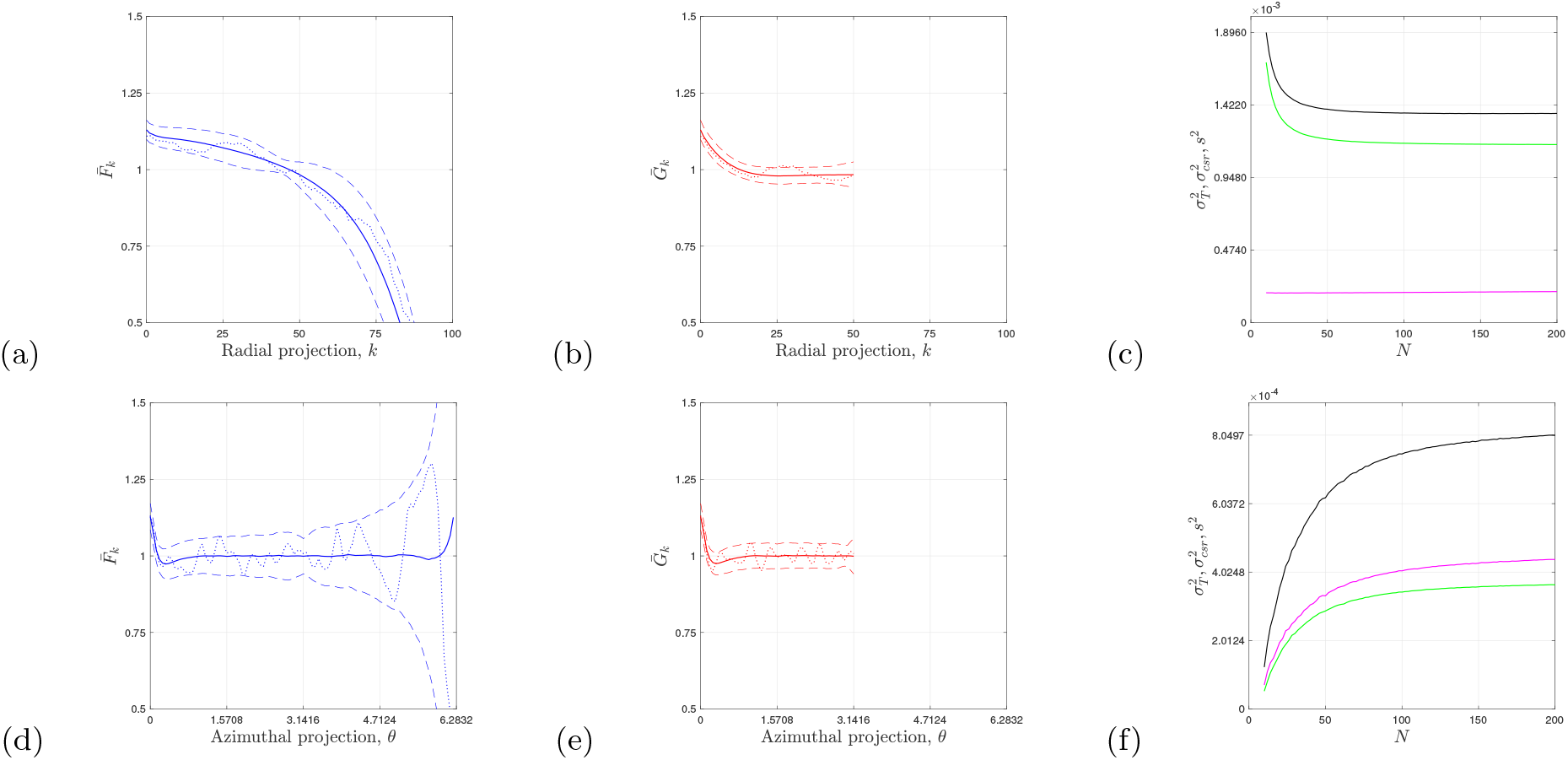
Diffusion-limited processes within a circular domain, *R* = 100, *D*_*I*_ = 1, *D*_*II*_ = *U*_*I*_ = *U*_*II*_ = 0, *δt* = 1, *δr* = 2. Initially, *n*^*I*^ (0) = 4000 agents of species *I* were distributed uniformly at random throughout the circle with a single agent from species *II* located at the origin with *n*^*II*^ (0) = 1. The simulations were terminated when all the agents in species *I* had transformed into species *II*. (a), (b), (d), (e) Average of PCFs calculated from *M* = 1000 realisations, solid curves. The dashed curves are for 2 standard deviations from the mean. The dotted curves are for a single realisation. Non-periodic (blue) and periodic (red) PCFs, *N* = 100. (c) and (f) Analysis of variance from *M* = 1000 realisations, 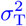 (black), 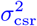 (green), *s*^2^ (magenta). (a)-(c) Projection in *r*-direction. (d)-(f) Projection in *θ*-direction.

**FIG. 8.**
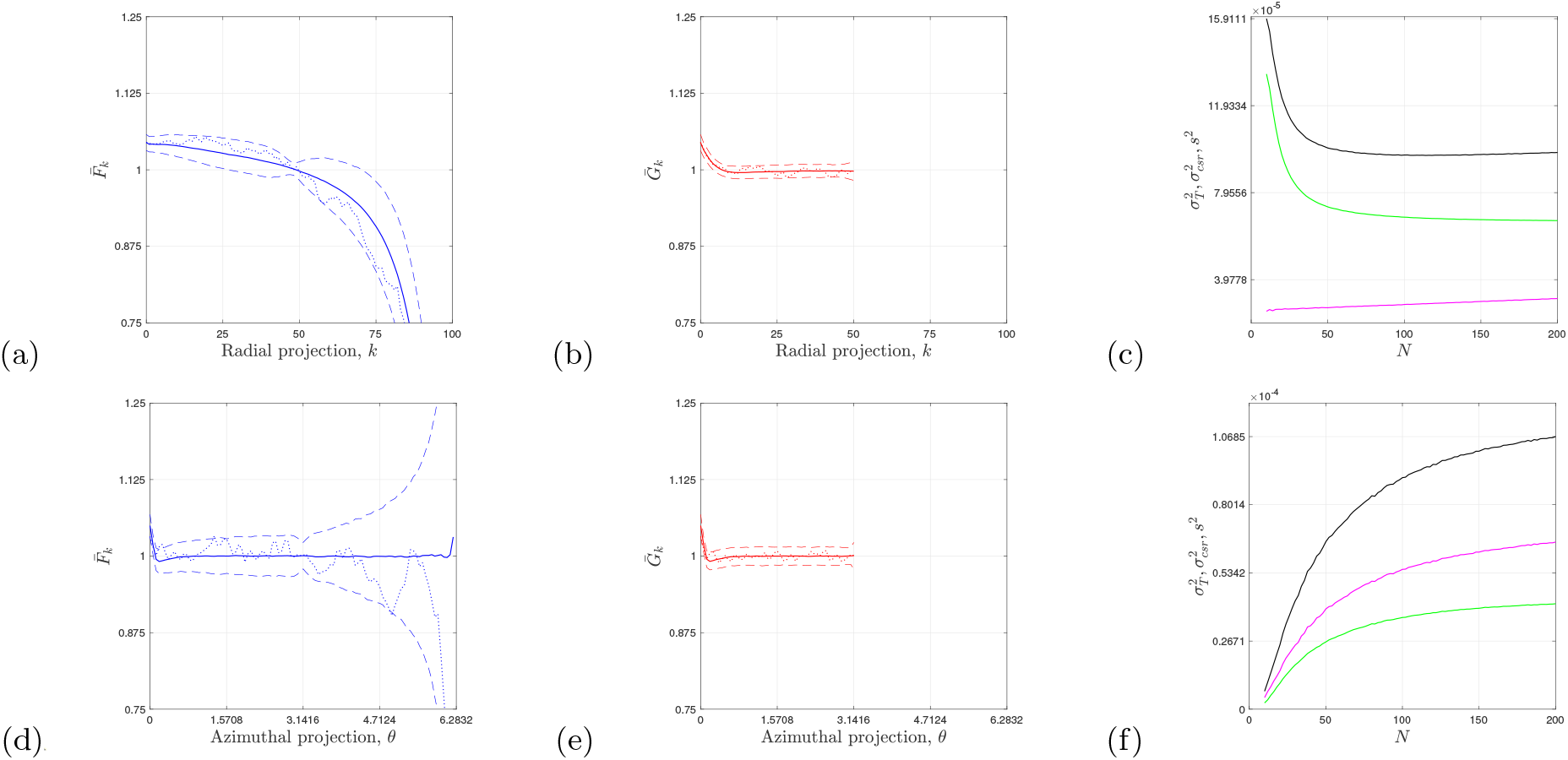
Diffusion-limited processes within a circular domain, *R* = 100, *D*_*I*_ = 1, *D*_*II*_ = *U*_*I*_ = *U*_*II*_ = 0, *δt* = 1, *δr* = 2. Initially, *n*^*I*^ (0) = 6000 agents of species *I* were distributed uniformly at random throughout the circle with a single agent from species *II* located at the origin with *n*^*II*^ (0) = 1. The simulations were terminated when all the agents in species *I* had transformed into species *II*. (a), (b), (d), (e) Average of PCFs calculated from *M* = 1000 realisations, solid curves. The dashed curves are for 2 standard deviations from the mean. The dotted curves are for a single realisation. Non-periodic (blue) and periodic (red) PCFs, *N* = 100. (c) and (f) Analysis of variance from *M* = 1000 realisations, 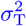 (black), 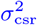 (green), *s*^2^ (magenta). (a)-(c) Projection in *r*-direction. (d)-(f) Projection in *θ*-direction.

We have observed that the PCFs can quantify aggregation and segregation at specific distances with reference to the CSR state. However, it is also valuable to measure a spatial pattern’s overall deviation from the reference CSR state [21–24]. To achieve this, we construct the weighted mean value of the PCF and the corresponding weighted variance. Along with the weighted mean value of sample variance (17), these measures enable us to determine both the departure of a pattern from the reference CSR state and the variability arising from the sampling of the DLP process. Additionally, the measures of variability introduced here provide a framework for quantifying the effects of varying the bin size *N* and the sample size *M* on the PCFs.

### A. Analysis of variance

An analysis of the variance provides a decomposition of the total variance, 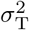, into a component for the sampling process, *s*^2^ (17), and a component for the processes departure from the CSR state, 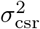. In this work, we derive expressions for the periodic PCF (with an even number of bins, *N*) and note that similar expressions hold in the non-periodic case. We then use the analysis of variance in the periodic case to characterise the aforementioned decomposition of variability for DLP processes.

For even *N*, we begin by defining the weighted mean-value of the periodic PCF (14) as

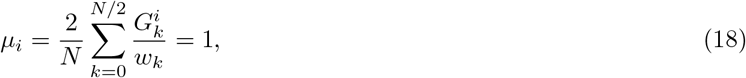

which by definition is constant and equal to unity for any realisation. The corresponding weighted variance is then defined as

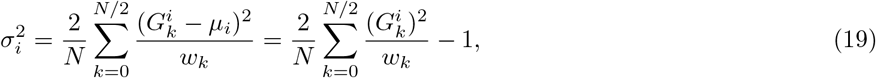

which is an overall measure of the *i*th realisation of the DLP process from the CSR state. The *grand* average of the weighted mean (18) and weighted variance (19) from a total of *M* realisations is then given by

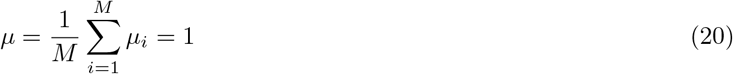

and

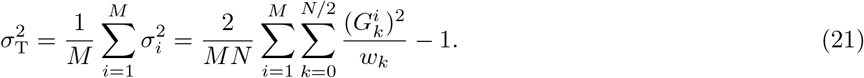

Equation (21) is the grand total measure of the variability. In addition, we also consider the weighted mean-value of the average PCF, and using (16) and (18) we find

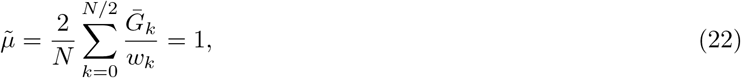

with the corresponding weighted variance

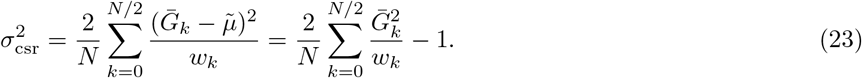

The variability (23) is a measure of the processes departure from the CSR state. The final measure of variability is the weighted mean of the variance in (17)

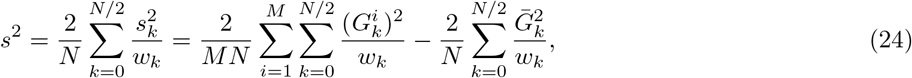

which is measure of the overall variability in the sample of realisations of the process.

The three measures of variability (21, 23, 24) are related by the expression

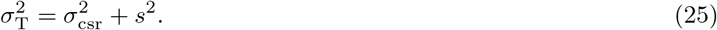

Equation (25) shows that the total variability 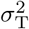 can be decomposed into a component representing the spatial deviation from the CSR state 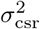 and a component for the variability across the realisations of the process *s*^2^.

It is interesting to consider some idealised scenarios and the corresponding values of the three measures of variability. When 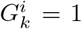 for all *i, k*, we have 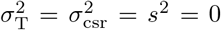 realisation is exactly at the CSR state with no variability across realisations. In practice, however, this is very unlikely to occur, and a CSR limit will be reached instead [23, 24]. Another idealised scenario is the most segregated state, where all agents reside in a single bin. In this case, (12) gives *c*_0_ = *n*^*j*^(*n*^*j*^ − 1)*/*2, with 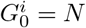 and 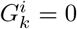 for *k*≠0 in (14). Furthermore, if this is the *same* single bin across all realisations, we have the maximum 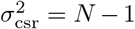, and, as previously, no variability across realisations, with *s*^2^ = 0. This naturally leads us to define the indices of variability

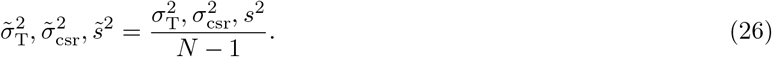

We now use the three measures (25) to quantify the variability of the DLP and to examine the effect of varying the number of bins, *N*, for the three different conditions, *n*^*I*^ (0) = 2000, 4000, 6000, shown in Figs. 6, 7, and 8 (third column).

For the radial projection, with the condition *n*^*I*^ (0) = 2000, we observe that the majority of the variability can be attributed to the DLP deviation from the CSR state, with much smaller variation due to the sampling of the process, as shown in Fig. 6(c). This is consistent with the results in (b), where the average periodic PCF signal (with *N* = 100 bins) is not close to unity for most values of the radial distance, with relatively compact confidence intervals. However, for the azimuthal projection, we find that the variation is split approximately in half, with both variability measures contributing more or less equally to the total variability [Fig. 6(f)]. These observations are also consistent with the results for the azimuthal projection in (e), where we see significant aggregation at short distances in the signal (i.e. deviation from the CSR state) and larger fluctuations around unity for moderate distances.

A qualitatively similar assessment holds in the radial case for the conditions *n*^*I*^ (0) = 4000, 6000, as shown in Figs. 7(c) and 8(c). In contrast, there is a change in the qualitative behaviour for the azimuthal projection, where the variability is no longer split evenly, with most of the variability being due to the sampling of the process instead of the deviation of the process from the CSR state [Figs. 7(f) and 8(f)].

For the condition with *n*^*I*^ (0) = 4000, we examine the effect of varying the number of realisations of the DLP for a fixed number of bins in Fig. 9. Using the indices of variability (26), we see that as few as 100 realisations produce results similar to those from 1000 simulations, for both projections.

**FIG. 9.**
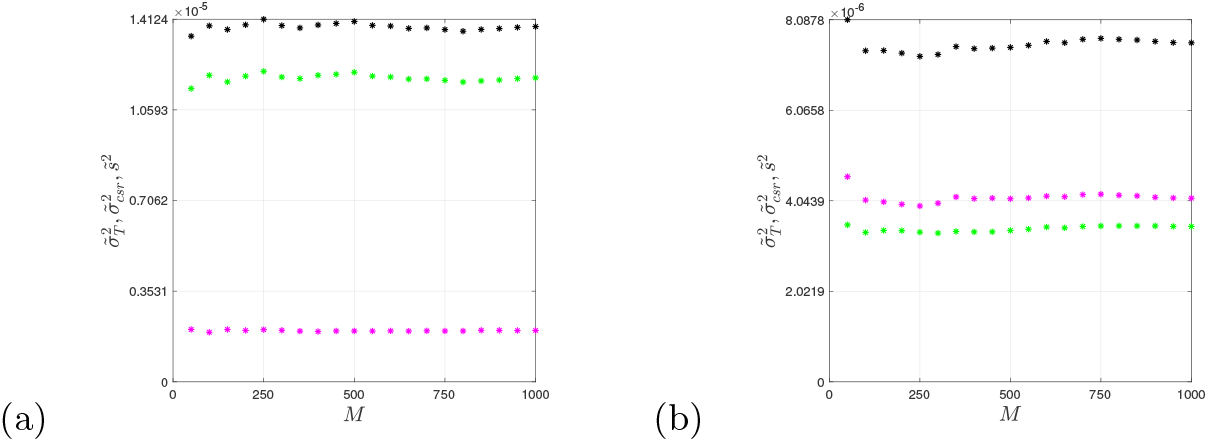
Diffusion-limited processes within a circular domain, *R* = 100, *D*_*I*_ = 1, *D*_*II*_ = *U*_*I*_ = *U*_*II*_ = 0, *δt* = 1, *δr* = 2. Initially, *n*^*I*^ (0) = 4000 agents of species *I* were distributed uniformly at random throughout the circle with a single agent from species *II* located at the origin with *n*^*II*^ (0) = 1. The simulations were terminated when all the agents in species *I* had transformed into species *II*. Analysis of variance with a fixed number of bins, 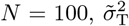 (black), 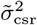 (green), 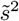 (magenta). (a) Projection in *r*-direction. (b) Projection in *θ*-direction.

We conclude our analysis of DLP in the 2D (*r, θ*)-plane with the condition *n*^*I*^ (0) = 4000 and two different values of the diffusion coefficient, *D*_*I*_ = 0.25 and *D*_*I*_ = 1. A realisation of the DLP with a value of *D*_*I*_ = 0.25 is presented in Fig. 10(a), which is similar to the realisation in Fig. 5(d), for a value *D*_*I*_ = 1. Importantly, the average periodic PCFs can distinguish the two processes with different rates of diffusion in both projections [Fig. 10(b,d)]. Interestingly, and in the case when *D*_*I*_ = 0.25, we observe that when 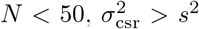 and when 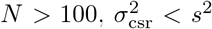 for the radial projection [solid curves, Fig. 10(e)]. This suggests some care needs to be taken when selecting the number of bins in the evaluation of the PCFs, and a spectral analysis of PCF bin-width [32] has shown to provide insight into this choice.

**FIG. 10.**
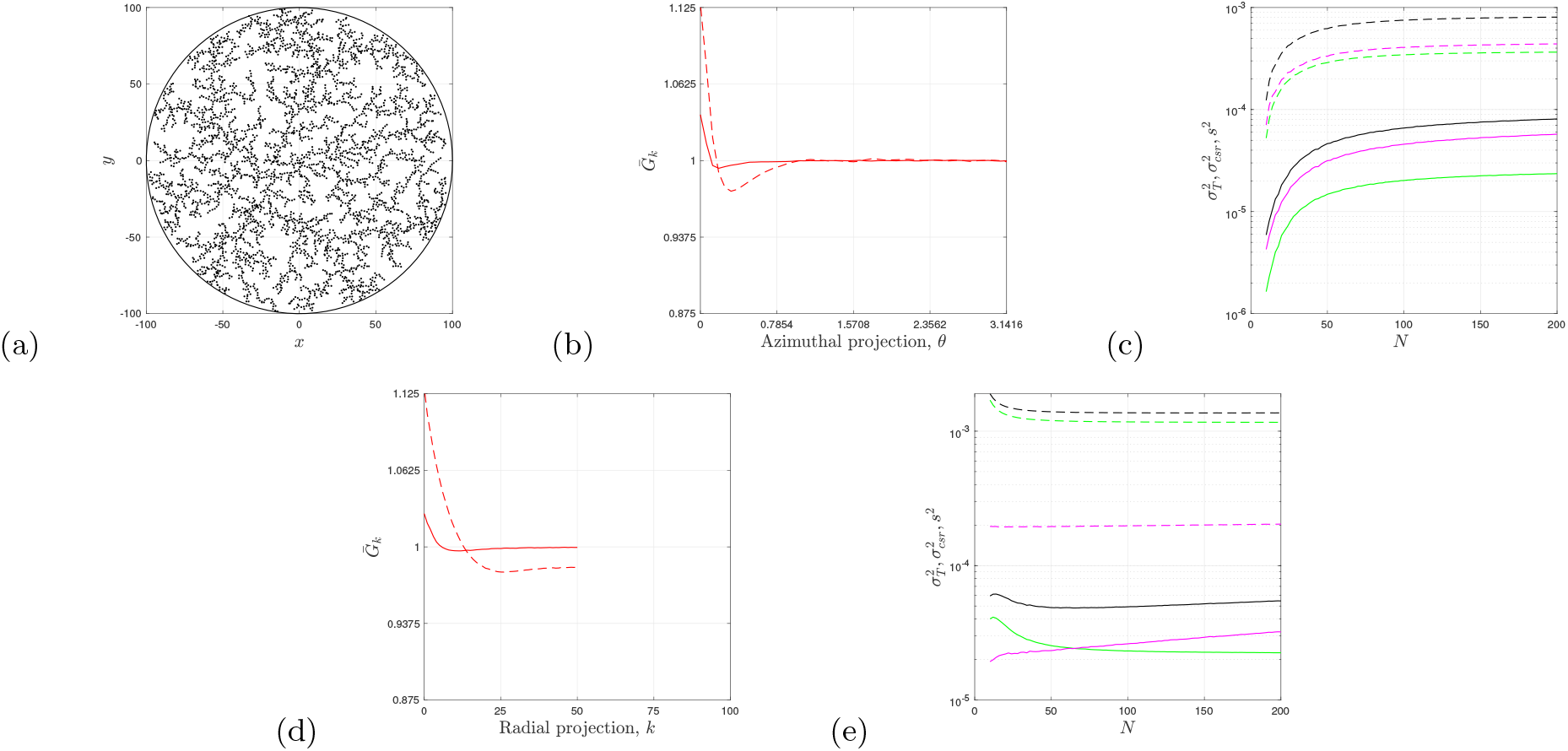
Diffusion-limited processes within a circular domain, *R* = 100, *D*_*II*_ = *U*_*I*_ = *U*_*II*_ = 0, *δt* = 1, *δr* = 2. Initially, *n*^*I*^ (0) = 4000 agents of species *I* were distributed uniformly at random throughout the circle with a single agent from species *II* located at the origin with *n*^*II*^ (0) = 1. The simulations were terminated when all the agents in species *I* had transformed into species *II*. (a) Realisation, *D*_*I*_ = 0.25, *t* = 1245. (b) and (d) Average periodic PCFs with *D*_*I*_ = 0.25 (solid curves) and *D*_*I*_ = 1 (broken curves), *M* = 1000. (c) and (e) Analysis of variance in azimuthal and radial direction, 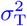 (black), 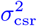 (green), *s*^2^ (magenta). Solid curves, *D*_*I*_ = 0.25. Broken curves, *D*_*I*_ = 1.

In this work, we also provide a spectral analysis of the periodic PCFs, deriving a relationship between the average periodic PCF and the coefficients of a discrete cosine transform. As we will demonstrate, the variability measure 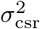 can be expressed in terms of these coefficients, and an analysis of the power spectrum can reveal characteristic distances in the DLP. In the Cartesian and azimuthal projections with equal bin widths, the characteristic dimensionless distances are directly related to corresponding characteristic physical distances.

### B. Spectral analysis

The discrete cosine transform [9, 32, 33], for even *N*, of the average periodic PCF is defined by

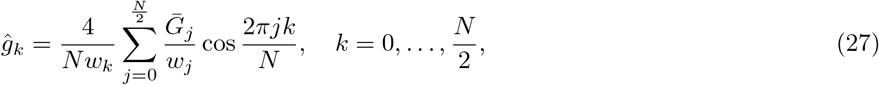

with inverse transform

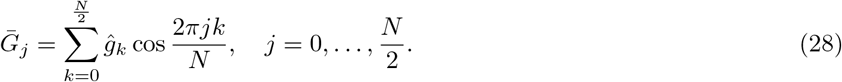

The coefficients are also related to the mean-value and variance of the average periodic PCF, and after some algebra involving Eqs. (16), (22), (23), (27), and (28), we arrive at the following expressions that relate the average periodic PCF (and its mean-value and variance) to the transform coefficients 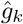

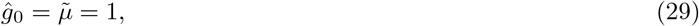

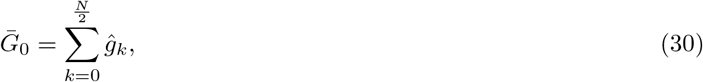

and

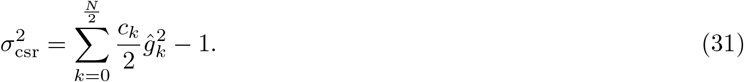

When plotting the power spectrum, 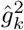, we exclude the constant and fixed value 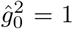 Coefficient. Examining the power spectrum can provide a systematic approach to identify dominant modes and their corresponding wavelengths in DLP.

In the remainder of this paper, we employ the average periodic PCF to analyse 3D DLP patterns within a cylindrical domain. Our first set of results focuses on six initial seed agents from species *II*, which are positioned on the interior of the cylinder’s surface. These seed agents are evenly spaced in both the *θ* and *z* directions and grow through DLP, while the domain is uniformly populated at random with three different initial densities of species *I, n*^*I*^ (0) = 2000, 4000, 6000. The simulations terminate when all agents of species *I* have fully transformed into species *II*, as depicted in the first row of Fig. 11. Given this setup, we expect the PCFs to effectively capture the regular patterning along both the *θ* and *z* directions.

**FIG. 11.**
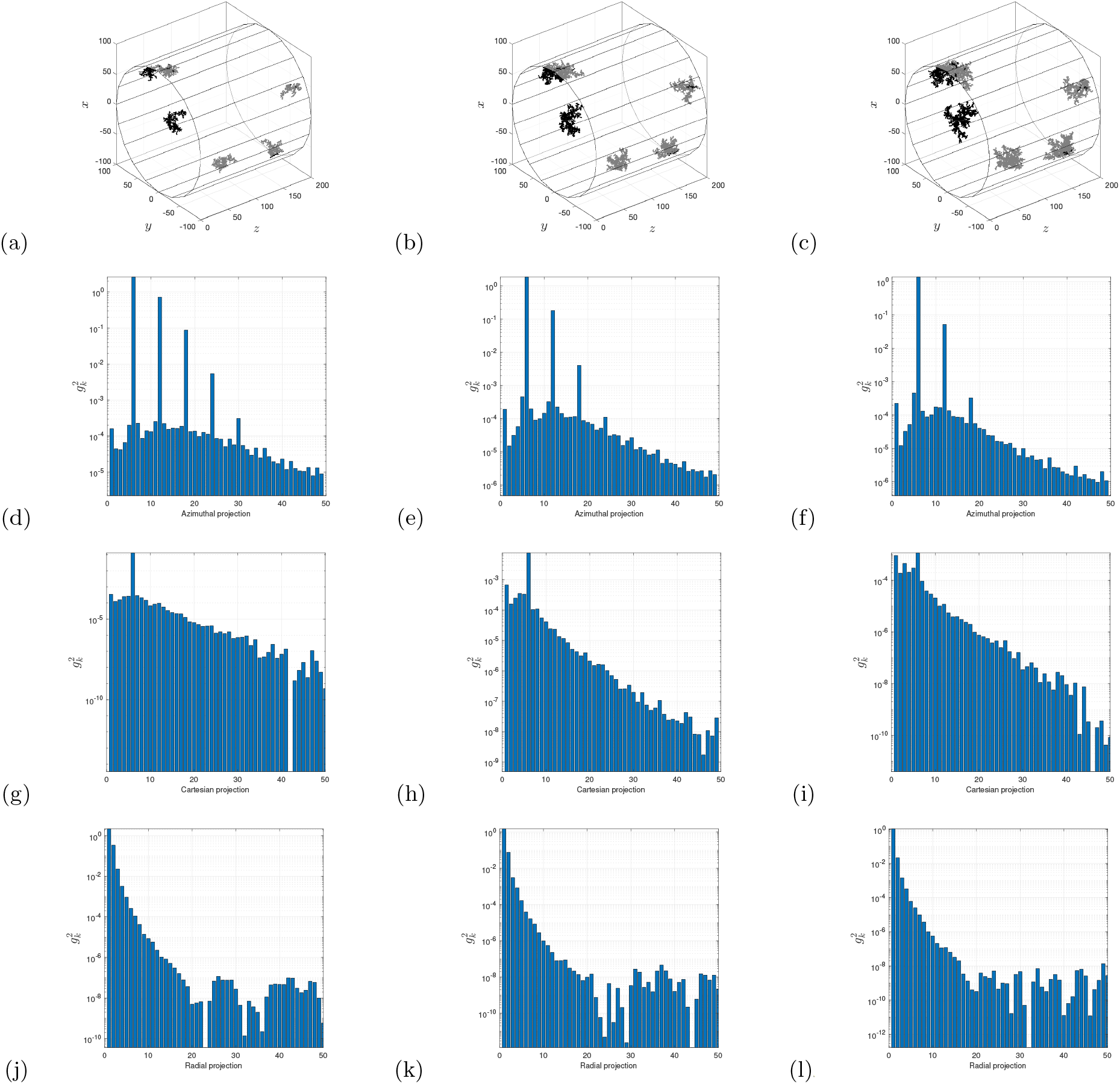
Diffusion-limited processes within a cylindrical domain, *R* = 100, *Z* = 200, *D*_*I*_ = 1, *D*_*II*_ = *U*_*I*_ = *U*_*II*_ = 0, *δt* = 1, *δr* = 2, *N* = 100, *M* = 100. Initially, six agents from species *II* were placed at 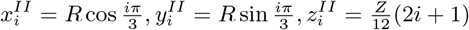 for *i* = 0, …, 5, with *n*^*II*^ (0) = 6. First to third column, initially *n*^*I*^ (0) = 2000, 4000, 6000 agents of species *I* were distributed uniformly at random throughout the cylinder. The simulations were terminated when all the agents in species *I* had transformed into species *II*. (a)–(c) Single realisations. Second to fourth row, power spectrums for the average periodic PCFs (see Fig. 12), excluding the 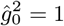 coefficient. (d) – (f) Azimuthal projection. (g)–(i) Cartesian projection. (j)–(l) Radial projection.

The power spectrum, 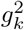 of the average periodic PCFs for each of the three initial setups (*n*^*I*^ (0) = 2000, 4000, 6000) is shown in the corresponding column beneath a typical realisation in the first row of Fig. 11. The second to fourth rows display the power spectrums for the azimuthal, Cartesian, and radial projections, respectively. In the azimuthal projection (second row), we observe that the coefficient with the largest magnitude corresponds to *k* = 6, yielding a characteristic length of *λ* = *L/k* = *π/*3 in the azimuthal direction. This precisely matches the angular spacing of the six initial seed agents of species *II*. For simulations with smaller aggregates (first column), we observe harmonics at higher frequencies due to the six clusters being almost completely separated in the azimuthal direction. In contrast, for larger clusters [(Fig. 11)(e) and (f)], the fundamental mode remains dominant, but the higher-frequency harmonics become less pronounced relative to neighbouring coefficients in the spectrum. As cluster size increases, greater overlap among the six clusters in the azimuthal direction results in a reduction in the magnitude of the higher-frequency modes. Thus, the spectrum successfully captures the expected fundamental length scale across all simulations, while also producing distinct and easily distinguishable power spectrums for the three different setups.

Similar to the analysis in the azimuthal direction, we find that the power spectrum effectively captures the characteristic length scale of 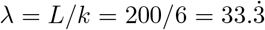 in the Cartesian direction (third row, Fig. 11), which corresponds to the spacing of the six initial seed agents of species *II*. However, in this direction, the clusters remain connected for all three initial setups (*n*^*I*^ (0) = 2000, 4000, 6000), and no visually identifiable higher-frequency modes are present. In contrast, and as expected, no dominant mode emerges in the radial direction (fourth row, Fig. 11).

The average periodic PCFs for each of the three setups (*n*^*I*^ (0) = 2000, 4000, 6000) are shown in Fig. 12. In both the azimuthal and Cartesian directions, we observe the fundamental frequencies, with a decrease in amplitude as cluster size increases due to the greater overlap in these directions. In the radial direction, the signal reaches a maximum for the setup with the largest clusters, indicating aggregation at short radial distances. Notably, the signals remain distinct across all three directions.

**FIG. 12.**
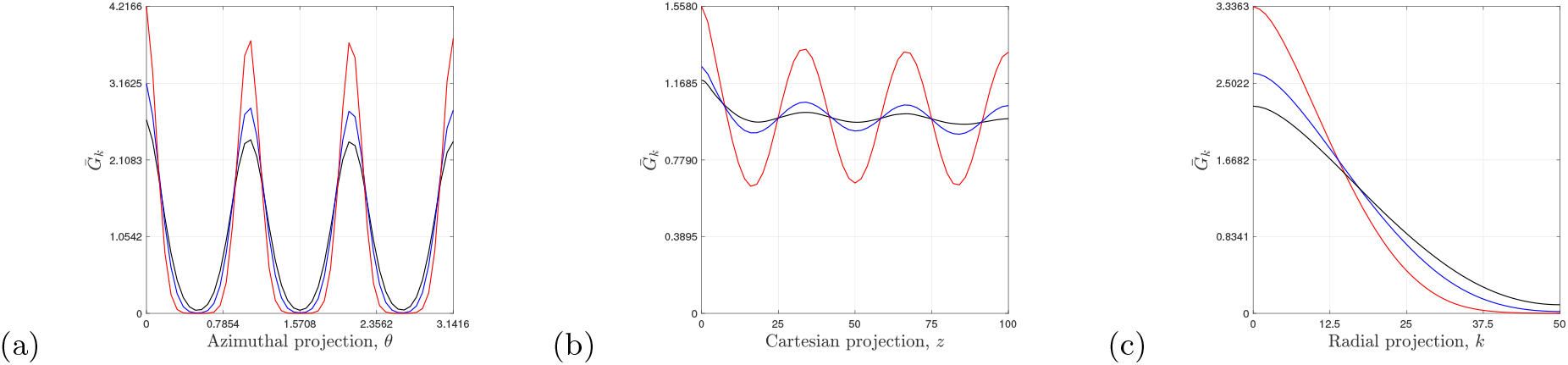
Average periodic PCFs for the results shown in Fig. 11, *n*^*I*^ (0) = 2000 (red), *n*^*I*^ (0) = 4000 (blue), *n*^*I*^ (0) = 6000 (black). (a) Azimuthal projection. (b) Cartesian projection. (c) Radial projection.

The results for the regular spacing of the six seed agents (Figs. 11 and 12) are compared to those for six seed agents placed uniformly at random along the interior surface of the cylinder (Fig. 13). Individual realisations for the random placement of seed agents look similar to those with regular spacing of seed agents (see first row of Fig. 11). The corresponding power spectrums and average periodic PCFs for *n*^*I*^ (0) = 2000 are presented in the first and second rows of Fig. 13, respectively. Unlike the regularly spaced case, where the PCFs exhibit oscillatory behaviour at moderate to large distances, as shown by the red curves in Figs. 12(a) and (b), no dominant mode is observed in either the azimuthal or Cartesian directions, see power spectrums in Figs. 13(a) and (b). Instead, for randomly placed seed agents, short-range aggregation is evident, with the signals flattening out to nearly constant values at larger distances [Figs. 13(d) and (e)]. This flattening indicates a lack of spatial structure, consistent with the random initial placement of seed agents, while short-distance aggregation corresponds to the increase in species *II* agents within each of the six (randomly distributed) clusters. As before, the signals in Fig. 13 differ significantly from those in Fig. 12 (red curves), further demonstrating the ability of PCFs to distinguish between spatial patterns that may appear visually similar at first glance.

**FIG. 13.**
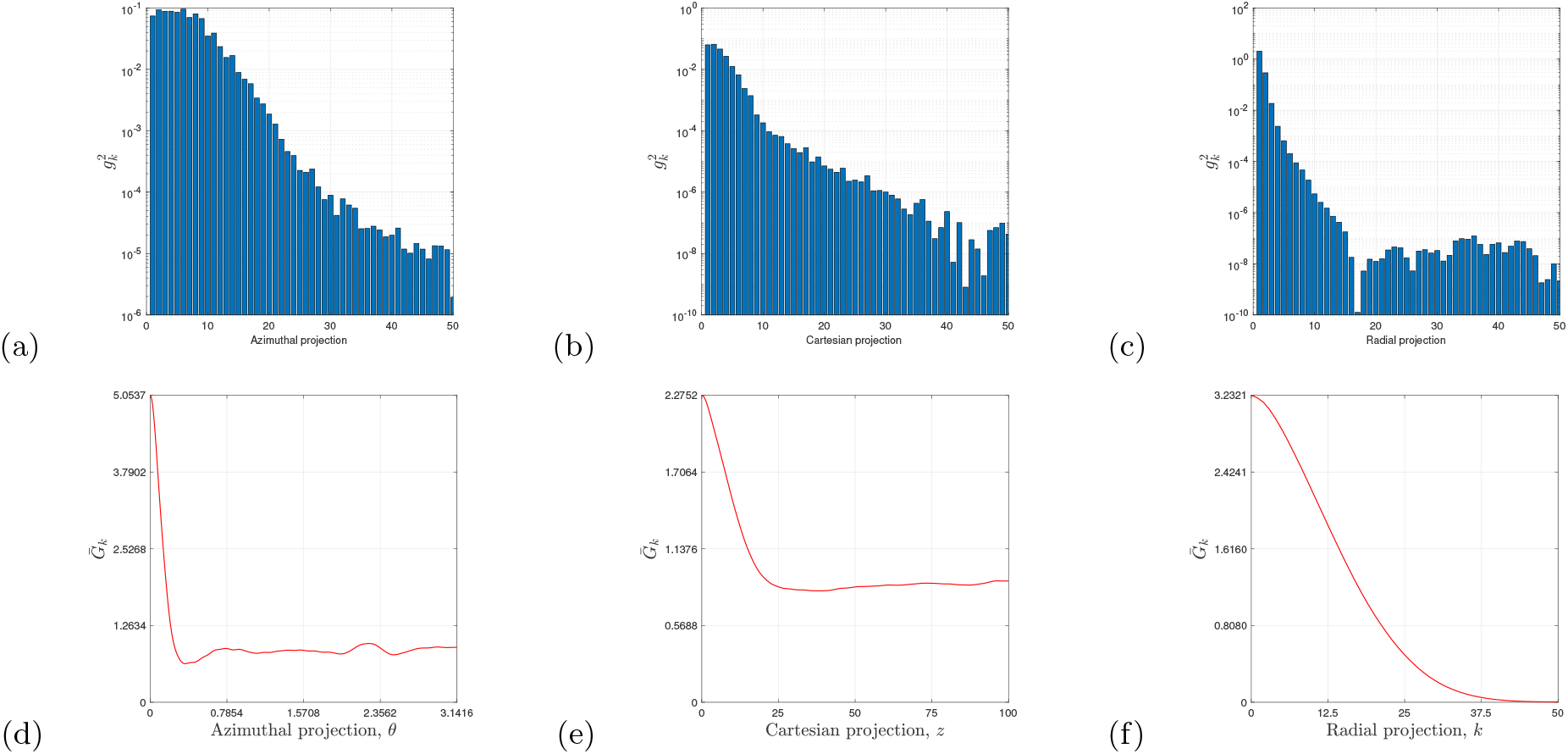
Diffusion-limited processes within a cylindrical domain, *R* = 100, *Z* = 200, *D*_*I*_ = 1, *D*_*II*_ = *U*_*I*_ = *U*_*II*_ = 0, *δt* = 1, *δr* = 2, *N* = 100, *M* = 100. Initially, six agents from species *II* were placed uniformly at random along the surface of the cylinder with *n*^*II*^ (0) = 6. Initially *n*^*I*^ (0) = 2000 agents of species *I* were distributed uniformly at random throughout the cylinder. The simulations were terminated when all the agents in species *I* had transformed into species *II*. (a)–(c) Power spectrums for the average periodic PCFs, excluding the 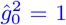 coefficient. (d)–(f). Average periodic PCFs. (a,d) Azimuthal projection. (b,e) Cartesian projection. (c,f) Radial projection.

While we have demonstrated that PCFs can identify characteristic length scales for a given process, we now turn our attention to variations in the DLP which are summarised in Tables I and II. Table I presents the values of 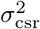, allowing us to assess pattern variation relative to the CSR state. The first three columns correspond to results obtained with regularly spaced seed agents (Figs. 11 and 12), whereas the final column (highlighted in blue) represents results for randomly placed seed agents (Fig. 13).

**TABLE I.** Departure from CSR state as measured by 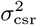, for the results shown in Figs. 11, 12 and 13. First three columns: Initial set-up with regular spaced seeds agents as specified in the caption of Fig. 11. Last column: Initial set-up with random placement of seed agents as specified in the caption of Fig. 13.

| $\sigma_{\text{csr}}^2$ | $n^I(0) = 2000$ | $n^I(0) = 4000$ | $n^I(0) = 6000$ | $n^I(0) = 2000$ |
| --- | --- | --- | --- | --- |
| Azimuthal projection | 1.7124 | 1.0244 | 0.7115 | 0.4535 |
| Cartesian projection | 0.0645 | 0.0048 | 0.0017 | 0.1124 |
| Radial projection | 1.2557 | 0.7839 | 0.5406 | 1.1835 |

For the regularly spaced seed agents (Figs. 11 and 12), we observe a decrease in 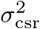 as *n*^*I*^ (0) increases across all three projections (see the first three columns of Table I). This trend is intuitive as a higher number of *n*^*I*^ (0) introduces greater random spatial variability around the location of the seed agents, which in turn smooths out the regular patterning imposed by their initial placement.

One might then expect lower values of 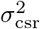 for the randomly placed seed agents compared to the regularly spaced case, given the same initial value of *n*^*I*^ (0) = 2000 (see the first and fourth columns in Table I). Indeed, this intuition holds in the azimuthal projection. However, in the Cartesian direction, we observe a counterintuitive result where the variation 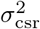 is actually larger for the randomly placed seed agents.

This unexpected behaviour can be partially explained by revisiting the power spectrums in Fig. 11. In the Cartesian projection, the dominant mode is less pronounced compared to the azimuthal projection. A similar trend is evident in the interpolants of Fig. 12, where the amplitude of oscillations is much greater in the azimuthal direction, indicating that clusters remain more segregated in this direction. In contrast, as previously noted, there is significant overlap in clustering in the Cartesian projection, leading to smaller oscillation amplitudes, and less segregation of the clustering in this direction. Consequently, the increased overlap in the Cartesian direction for the regularly placed seed agents results in less overall variation from the reference CSR state than the randomly placed seed agents.

A similar trend is observed in the variation due to sampling for the regularly spaced seed agents, as *n*^*I*^ (0) increases, *s*^2^ decreases (see the first three columns in Table II). This result is expected, as larger values of *n*^*I*^ (0) smooth out the variation in individual realisations, reducing overall variability in the sampling process.

**TABLE II.** Variability in realisations measured by *s*^2^, for the results shown in Figs. 11, 12 and 13. First three columns (black): Initial set-up with regular spaced seeds agents as specified in caption of Fig. 11. Last column: Initial set-up with random placement of seed agents as specified in caption of Fig. 13.

| $s^2$ | $n^I(0) = 2000$ | $n^I(0) = 4000$ | $n^I(0) = 6000$ | $n^I(0) = 2000$ |
| --- | --- | --- | --- | --- |
| Azimuthal projection | 0.0149 | 0.0091 | 0.0069 | 0.3737 |
| Cartesian projection | 0.0056 | 0.0018 | 0.0010 | 0.1004 |
| Radial projection | 0.0106 | 0.0067 | 0.0061 | 0.0134 |

When comparing the results for regular and random seed placement with *n*^*I*^ (0) = 2000 (see the first and fourth columns in Table II), the trend in variation is more intuitive than before with larger values of *s*^2^ being observed for the randomly placed seed agents. This outcome aligns with our expectations, as random placement introduces greater variability across different realisations in the sampling process.

In the final set of results (Fig. 14), we again examine 3D DLP within a cylindrical domain, now incorporating a nonzero and spatially variable drift

**FIG. 14.**
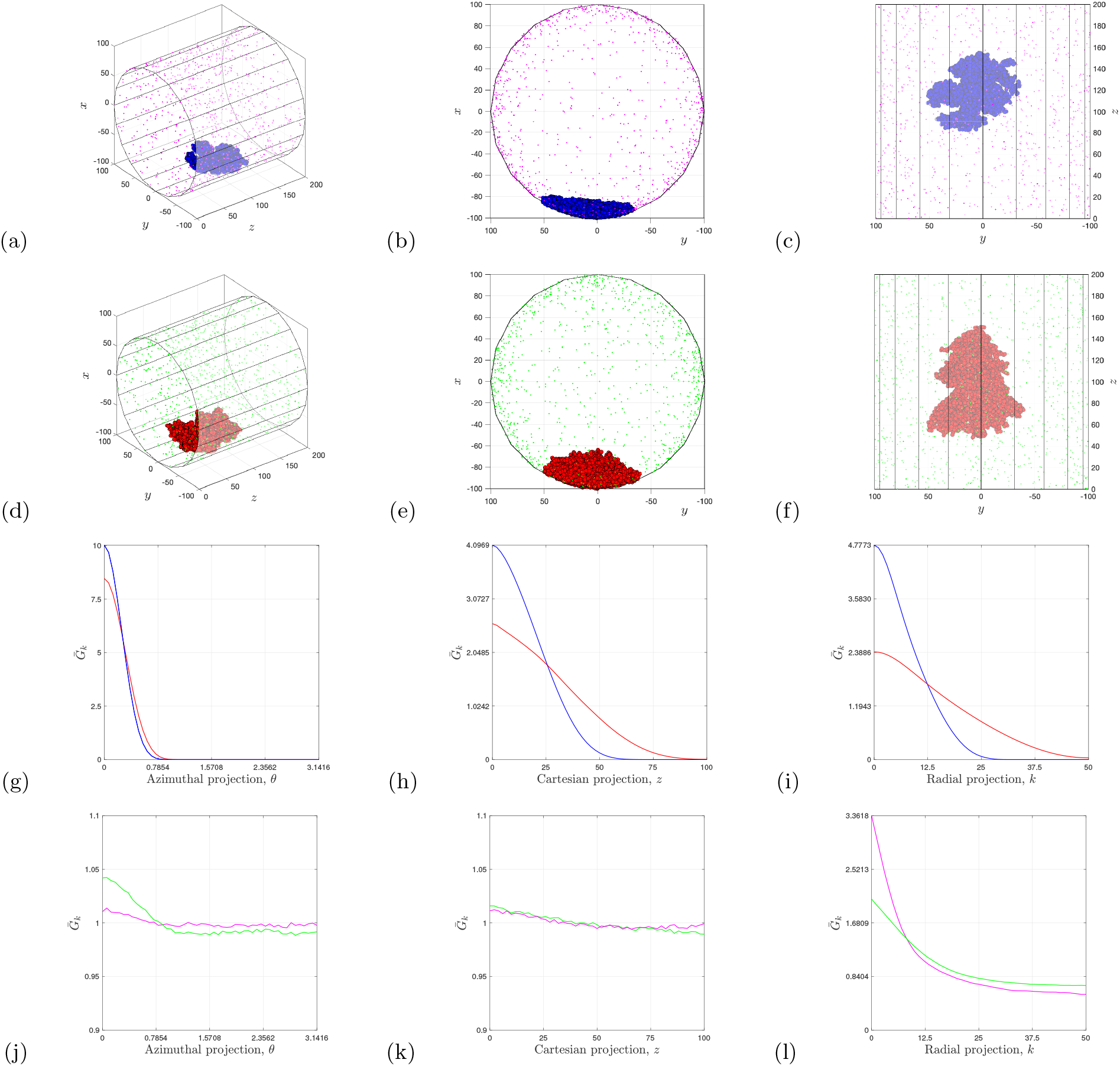
Diffusion-limited processes within a cylindrical domain, *R* = 100, *Z* = 200, *D*_*I*_ = 0.1, *D*_*II*_ = *U*_*II*_ = 0, *δt* = 1, *δr* = 2, *N* = 100, *M* = 100. (a)–(c) Simulation with drift *U*_*I*_ = 0.01. Species *I*, magenta. Species *II*, blue. (d)–(f) Simulation with drift *U*_*I*_ = 0.001. Species *I*, green. Species *II*, red. (g)–(l) Average periodic PCFs. (g)–(i) Comparison of species *II* for values of the drift *U*_*I*_ = 0.01 (blue curve) and *U*_*I*_ = 0.001 (red curve). (j) – (l) Comparison of species *I* for values of the drift *U*_*I*_ = 0.01 (magenta curve) and *U*_*I*_ = 0.001 (green curve). (g,j) Azimuthal projection. (h,k) Cartesian projection. (i,l) Radial projection.

**FIG. 15.**
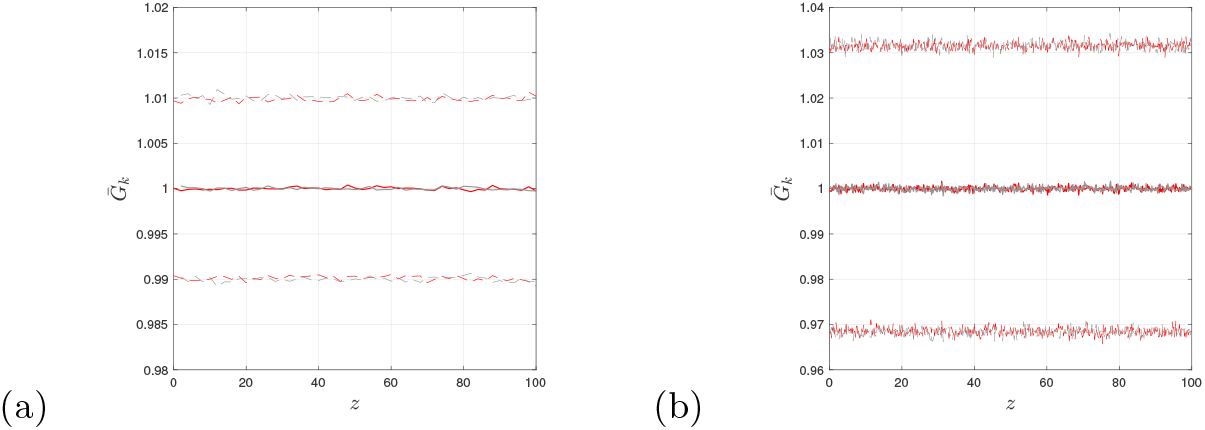
Cylinder uniformly populated with *n*^*I*^ = 2000 agents, *R* = 100, and *Z* = 200. The solid curves represent the average periodic PCF computed from *M* = 1000 realisations, for the Cartesian projection. Similar results are found for the other two projections (not shown). The dotted curves indicate two standard deviations from the mean. The red curves correspond to the formulation developed in this paper, based on pairwise combinations of bin counts. The grey curves correspond to the author’s previous formulation, which first computes pairwise combinations of individual agents before binning. (a) *N* = 100. (b) *N* = 1000.

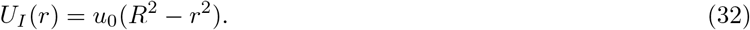

Poiseuille’s equation (32) describes a parabolic velocity profile for steady, unidirectional flow in the *z*-direction, with a no-slip condition at the cylinder’s interior surface and a maximum velocity along the centerline. Agents of species *I* are advected by the drift *U*_*I*_, while also undergoing additional random motion governed by the diffusion coefficient *D*_*I*_. The parameter *u*_0_ represents the ratio of the pressure drop per length of pipe to the viscosity of the base fluid.

At *t* = 0, the cylinder is initially populated with *n*^*I*^ (0) = 1000 agents of species *I*, distributed uniformly at random. A single seed agent of species *II* is placed on the interior surface of the cylinder at (− 100, 0, 150). During each timestep *δt*, the number of species *I* agents can decrease in two ways, either by transformation via DLP into species *II* or drift in the positive *z*-direction, exiting the cylinder at *z* = *Z*. To maintain a roughly constant population of species *I*, the decrease in agent number is replenished at *z* = 0, where new agents are introduced uniformly at random within the disc at *z* = 0. The simulations terminate when the total number of species *II* agents reaches *n*^*II*^ = 6000.

We present simulations for two different values of the drift with the same diffusion coefficient in the first two rows of Fig. 14. For the smaller value of *u*_0_, the aggregate or cluster of agents of species *II* (red) appears visually less dense compared to the aggregate (blue) observed for the larger *u*_0_. This suggests that increased drift enhances the concentration of agents within clusters, leading to more compact and localised pattern formation. Nonetheless, both simulations show directed growth towards the source of species *I* agents released in the disc at *z* = 0, which is a characteristic of DLP.

The average periodic PCFs for agent species *I* and *II*, for both values of *u*_0_, are shown in the third and fourth rows of Fig. 14, with PCF signal colours corresponding to the agent colours in each simulation. In the third row, the PCF signals for species *II* across all three projections indicate greater short-distance aggregation for the larger *u*_0_ value (blue), consistent with the formation of smaller, denser aggregates observed in the simulations (first two rows). Similarly, the quantified patterning of species *I* in the fourth row aligns with these findings. For instance, in the radial projection panel [Fig. 14(l)], increased short-distance aggregation is observed for the larger *u*_0_, which corresponds to the more concentrated clustering of magenta agents near the cylinder’s interior compared to the green agents, as seen in Fig. 14(b) and (e).

These results further highlight the effectiveness of PCFs in quantifying multi-species patterning, offering a robust framework for analysing spatial correlations in more complex models of diffusion-limited processes.

## IV. DISCUSSION

In this study, new one-dimensional non-periodic and periodic pair-correlation functions (PCFs) have been developed to analyse the projected patterns generated by diffusion-limited processes (DLP) within a cylindrical spatial domain. These PCFs build upon the author’s previous work on one-dimensional PCFs but with a key refinement. In the prior formulation, PCFs were computed by evaluating the distances between all pairwise combinations of individual agents, followed by binning and normalisation with respect to the complete spatial randomness (CSR) state. However, this approach becomes computationally infeasible when dealing with large populations due to the number of combinations in pairwise distance calculations. To overcome this limitation, the present work introduces a more computationally efficient approach. Instead of computing distances between all agent pairs directly, individual data points are first binned, and the PCF is then computed based on pairwise combinations of these bin counts, followed by normalisation with respect to the CSR state.

While one-dimensional PCFs have been valuable as summary statistics in advanced inference algorithms [36, 37], their high computational cost has often limited their practical use, particularly when millions of evaluations are required for parameter estimation. The approach introduced here is significantly more efficient, as the computational time remains independent of the number of individuals for a fixed number of bins. This enables faster and more scalable PCF evaluations, making them a more practical tool for parameter estimation in diffusion-limited systems.

In addition to enhancing computational efficiency, this study also introduces a straightforward analysis of the variability in PCF calculations—an aspect that has not been explored in previous studies on one-dimensional pair-correlation functions. The two components of this variability allow us to quantify both the global departure of a process from the CSR state and the variation due to sampling. Together, these analyses enable a comprehensive understanding of spatial patterns where PCF signals provide local information at specific distances within the domain, while global measures capture overall patterning and sampling effects. Furthermore, connecting PCF analysis with spectral methods offers additional insights into characteristic length scales and the extent of deviation from the CSR state.

The periodic PCF is found to be significantly more accurate than the non-periodic PCF, as demonstrated by the narrower 95 % confidence intervals in Figs. 4, 6, 7, and 8, consistent with previous studies on one-dimensional PCFs [25, 27, 32, 35]. Given this improved accuracy, we employ the periodic PCF to assess DLP patterning in Figs. 9–14. However, caution is required when interpreting periodic signals for inherently non-periodic problems or projections. The azimuthal projection is naturally periodic, making the periodic PCF an appropriate choice for analysing patterns in this direction. For the Cartesian projection, periodic PCFs are also suitable in Figs. 11–13, as we are effectively examining a section of a cylinder, where assuming periodic boundary conditions is both reasonable and useful for detecting periodic behaviour. In contrast, the radial projection is inherently non-periodic, making the non-periodic PCF generally more appropriate. However, in this study, DLP leads to clustering near the cylinder’s interior boundary, with only short-range aggregation and no correlations at moderate to large radial distances. Given this context, the more accurate periodic PCF remains a suitable choice for analysing DLP processes.

The DLP model in this study reproduced patterns reminiscent of the seminal work of Witten and Sander [16, 17] within a cylindrical framework [see Fig. 5]. Their work demonstrated that the density-density correlation function follows a fractional power law over distance within an aggregate or cluster, leading to scale-independent correlations over moderate to large distances. Our periodic PCF results for the azimuthal projection align with these findings, showing an approximately unit-valued signal at moderate and large distances [see Figs. 6–10]. This indicates the absence of a characteristic length scale beyond the short-range aggregation in this direction. The radial results further support this conclusion, particularly for patterns spanning the entire circular domain, where the signal flattens to a constant value [see Figs. 7–10]. The relationship between pair-distance functions (un-normalised PCFs) and the fractal dimension of DLP patterns remains an open question and is the subject of our ongoing research.

The 3D DLP modelling, combined with PCF analysis, highlights the potential applications of this method in future studies involving more complex models and multi-species interactions. As mentioned earlier, the model can be adapted to incorporate additional mechanisms, such as agent proliferation and removal (e.g. cell division and death in biological systems). Furthermore, integrating a velocity field to simulate fluid flow enables the study of physically significant processes, such as blood clot formation in blood vessels [38, 39], as well as microbial colony growth on the interior surfaces of tubes and the build-up on large-scale wastewater pipes, as illustrated in Fig. 1.

The PCFs developed in this study may also have applications in other areas, such as classification algorithms, which have been successfully used to characterize the morphology of dimorphic yeast growth, the structure of cancellous bone, and the marbling in beef [40, 41]. Additionally, the open-source image processing software TAMMiCol incorporates one-dimensional PCFs in its toolbox for automated image analysis. We plan to enhance this software by integrating the more efficient PCFs in the app [42]. Finally, the use of PCFs in moment closure models [43, 44] is another avenue for our future research.

## APPENDIX

### Accuracy and computational efficiency

A comparison of the periodic PCF computed using the bin-based approach developed in this paper and the pairwise agent-based approach from the author’s previous work shows that both methods achieve the same accuracy for 100 and 1000 bins. However, for 100 bins, the method presented in this paper required approximately 1 second to complete 1000 evaluations, whereas the previous method took about 30 seconds on a MacBook laptop. This represents a significant improvement in computational efficiency without any loss in accuracy.

### Spherical coordinates

The PCFs in this work can be adapted for other curvilinear coordinate systems, such as spherical coordinates, with corresponding position vectors

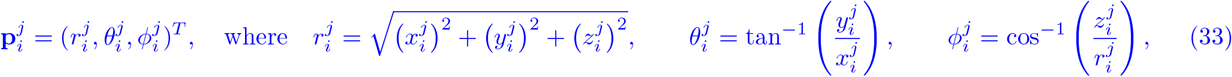

with 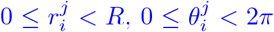 and 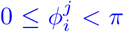. In the azimuthal direction, the bin-widths are constant and are the same as found in the cylindrical coordinate system (6). For the (spherical) radial case, we replace (7) on the interval [0, *L* = *R*] with

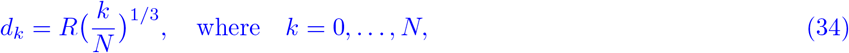

corresponding to equal volume spherical shells. For the polar projection on [0, *L* = *π*], we have the variable bin widths

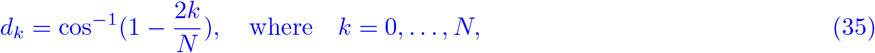

giving equal volume canonical shells. Similar results are found to the cylindrical case when the sphere is populated uniformly at random (not shown). Results for regular patterns within a sphere are presented in Fig. 16, thus demonstrating the application of our PCFs to analyse spatial patterns in other coordinate systems.

**FIG. 16.**
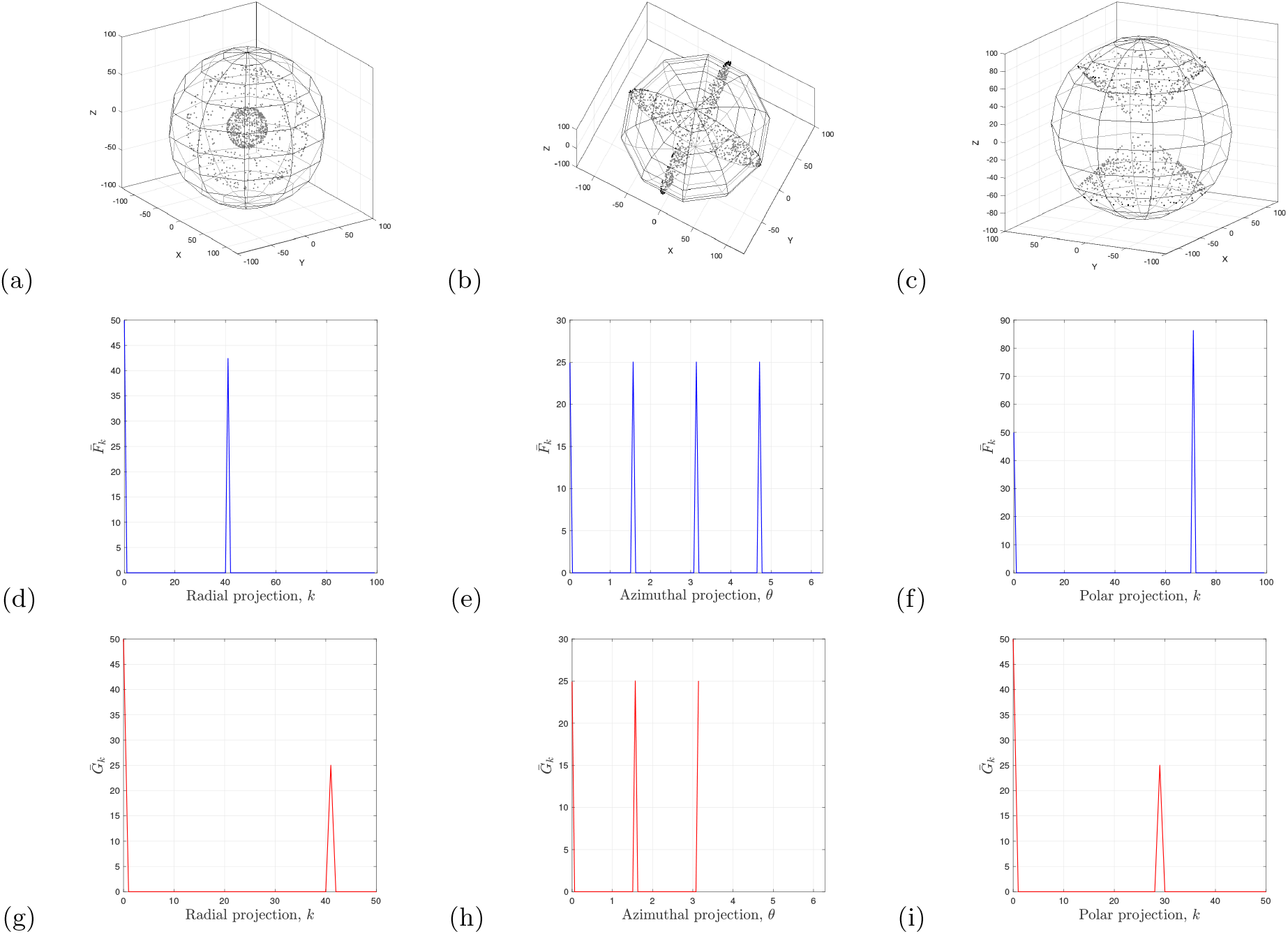
Regular spatial patterns within a sphere, *n*^*I*^ (0) = 1000 agents, *R* = 100, *N* = 100, *M* = 1000. (a) Regular pattern in radial direction with an even split of agents with radial distance 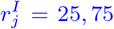 otherwise distributed uniformly at random. (b)Regular pattern in azimuthal direction with an even split of agents at angles 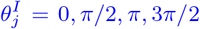, otherwise distributed uniformly at random. (c) Regular pattern in polar direction with an even split of agents at angles 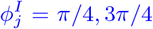, otherwise distributed uniformly at random. (d)–(f) Average PCF (blue). (g)–(i) Average periodic PCF (red). (d) and (g) Radial projection. (e) and (h) Azimuthal projection. (f) and (i) Polar projection.

## ACKNOWLEDGEMENTS

This work was supported by the Australian Research Council, DP230100406. We thank Natasha Binder for her assistance with this work.

